# Patterns of neural activity in prelimbic cortex neurons correlate with attentional behavior in the rodent continuous performance test

**DOI:** 10.1101/2024.07.26.605300

**Authors:** Jorge Miranda-Barrientos, Suhaas Adiraju, Jason J. Rehg, Michael S. Totty, Henry L. Hallock, Ye Li, Gregory V. Carr, Keri Martinowich

## Abstract

Sustained attention, the ability to focus on a stimulus or task over extended periods, is crucial for higher level cognition, and is impaired across multiple neuropsychiatric and neurodevelopmental disorders, including attention-deficit/hyperactivity disorder, schizophrenia, and depression. The rodent continuous performance test (rCPT) is a translational task that can be used to investigate the cellular mechanisms underlying sustained attention. Electrophysiological single unit and local field potential (LFPs) recordings reflect changes in neural activity in the prelimbic cortex (PrL) in mice performing sustained attention tasks. While evidence linking PrL neuronal activity to sustained attention is compelling, most studies have focused on single-cell activity surrounding behavioral responses, overlooking population-level dynamics across entire sessions that could offer additional insight into fluctuations in attention during task performance. Here, we used *in vivo* endoscopic calcium imaging to record patterns of neuronal activity in PrL neurons using the genetically encoded calcium sensor GCaMP6f in mice performing the rCPT at three timepoints characterized by differing levels of cognitive demand and task proficiency. We analyzed single-cell activity surrounding behavioral responses and population-level dynamics across sessions to determine how PrL neuronal activity evolves with sustained attention performance. A higher proportion of PrL neurons were recruited during correct responses in sessions requiring high task proficiency. Moreover, during rCPT sessions, mice intercalated responsive-engaged periods with non-responsive-disengaged periods. Applying a Hidden Markov Model (HMM) with two states to global PrL activity, we found neuronal states associated with task engagement. These states are characterized by different levels of correlated neuronal activity within PrL neurons. Overall, these findings illustrate that task proficiency, and task engagement differentially recruit activity in PrL neurons during a sustained attention task.

## Introduction

Sustained attention, the ability to focus on relevant stimuli or tasks over extended periods, is a fundamental component of cognitive processes [1]. Deficits in attention are frequently noted in individuals diagnosed with neuropsychiatric and neurodevelopmental disorders, including attention-deficit/hyperactivity disorder (ADHD) [2,3], schizophrenia [4,5,6], and depression [7,8]. Attention deficits are negatively correlated with functional outcomes and quality of life in these patients. Identifying the cellular and circuit mechanisms mediating sustained attention is a crucial step toward identifying novel pharmacological targets to treat attention deficits.

The most commonly used measure to quantify attention in neuropsychological batteries is the continuous performance test (CPT) [9]. The CPT has been used to assess attention deficits in patients with brain damage [10], schizophrenia [11] and ADHD [12]. A rodent CPT (rCPT) was developed as a touchscreen-based translational version of the human CPT, which requires mice to discriminate between a target stimulus (S+) and a non-rewarded stimuli (S-) in a series of successive trials [13,14]. The rCPT has predictive validity [13,14] as pharmacological agents that improve attention (e.g. amphetamine, methylphenidate, and atomoxetine) produce similar effects across species [6,15,16], and homologous brain regions are recruited during performance across species [6,15,16,17].

The human dorsal anterior cingulate cortex (dACC) is strongly linked to sustained attention. Brain imaging studies demonstrated dACC activation in subjects performing attentionally demanding tasks [18,19,20,21,22], including high-load tasks that incorporate presentation of distractors [23]. Moreover, dACC activity is decreased in individuals diagnosed with disorders that feature sustained attention deficits such as ADHD [3,24], schizophrenia [25,26], and obsessive-compulsive disorder [22]. In rodents, the prelimbic cortex (PrL) shows both functional and anatomical similarity to the human dACC [27,28]. Neuronal activity in the PrL is correlated with attention [17,29,30,31], while inhibiting neuronal activity in the PrL [32] or inputs to the PrL [33], disrupts sustained attention. Moreover, we reported changes in the power and coherence of PrL oscillations, as well as their directionality with locus coeruleus (LC) oscillations during rCPT performance [34]. However, how neuronal dynamics at the level of individual PrL neurons underlie attentional behavior remains to be elucidated.

In this study, we conducted *in vivo* endoscopic calcium imaging with a genetically encoded calcium sensor (GCaMP6f) to investigate neural dynamics in individual PrL neurons in mice performing the rCPT. We trained mice in the rCPT, and recorded calcium activity of PrL neurons at three time points (Stage 2, Stage 3-early, and Stage 3-late), which are characterized by differing levels of cognitive demand, and reflect different levels of task proficiency. We identified heterogeneous responses of PrL neurons during behavioral responses - specifically, a higher proportion of PrL neurons were modulated during correct responses, particularly in sessions requiring high levels of proficiency (Stage 3-late). Moreover, during rCPT sessions, mice intercalated responsive-engaged periods with non-responsive-disengaged periods. To test whether PrL neuronal activity reflects task engagement, we ran a Hidden Markov Model (HMM) with two states and found a higher probability of behavioral responses happening in one of the HMM identified states (Engaged state), while the probability of non-responsive periods was higher during the other state (Disengaged state). Importantly, neuronal activity during the disengaged state exhibited higher correlation across PrL neurons. Together, these results demonstrate that task engagement is linked to distinct neuronal states characterized by changes in network synchrony.

## Materials and Methods

### Animals

Male C57BL/6J mice (Strain #000664; The Jackson Laboratory, Bar Harbor, ME) were 8-10 weeks old at the start of the experiment. Mice were group housed (4/cage) in disposable polycarbonate caging (Innovive, San Diego, CA) and maintained on a reverse 12/12 light/dark cycle (lights on at 19:00 hours / lights off at 07:00 hours). Following surgeries for virus injection and gradient-index (GRIN) lens implantation, mice were single housed for the remainder of the experiment. Mice received Teklad Irradiated Global 16% Protein Rodent Diet (#2916; Envigo, Indianapolis, IN) in the home cage ad libitum until the start of the food restriction protocol, and water was available in the home cage ad libitum throughout all experiments. Behavioral testing was conducted Monday-Friday during the dark phase (07:00-19:00 hours). All experiments and procedures were approved by the Johns Hopkins Animal Care and Use Committee and in accordance with the Guide for the Care and Use of Laboratory Animals. Group size was selected based on our previous reports in the same task, and no randomization of animals was performed.

### Surgical procedures

Mice were anesthetized with isoflurane (induction: 2-4% in oxygen, maintenance: 1-2%) and secured to a stereotaxic frame. The top of the skull was exposed by an incision along the midline of the scalp, Bregma and Lambda were identified, and the head was leveled to ensure the skull was flat. A small hole was drilled with a 0.9 mm burr (Fine Science Tools, Foster City, CA) above the PrL, and 400 nl of a viral vector encoding the fluorescent calcium sensor (AAV1.Syn.Flex.GCaMP6f.WPRE.SV40; titer ≥ 1×10¹³ vg/mL) was injected into the PrL (AP: +1.8; ML: ±0.3, DV: −1.4). Injections were made using a Micro4 controller and UltraMicroPump along with a 10 µl Nanofil syringes equipped with 33-gauge needles (WPI Inc., Sarasota, FL). The syringe was left in place for 10 minutes after injection to minimize diffusion. Following viral injections, a gradient-index (GRIN) lens was implanted directly above the PrL. First, a slightly larger hole was drilled with a 1.8 mm burr (Fine Science Tools, Foster City, CA) at the viral injection site. Blood was cleaned using a sterile saline solution and swabbed until bleeding stopped, and the skull hole was clear. Then, a 4.00 mm X 1.00 mm GRIN lens integrated with a baseplate (Inscopix Inc., San Diego, CA) was slowly lowered using the stereotaxic frame at a rate of 0.2 mm / min until it reached 300 µm above the PrL (GRIN lens coordinates: AP: 1.7, MV: ±0.3, DV: −1.5). The GRIN lens was then secured to the skull using three small skull screws Fine Science Tools, Foster City, CA) positioned along the skull for extra support, and black dental acrylic (Ortho-jet, Lang Dental manufacturing Co., Wheeling, IL) to obscure outside light. Following surgery, the incision site not covered by the dental acrylic was closed using surgical staples (Fine Science Tools, Foster City, CA), and animals recovered on a heating pad for 30-60 mins before being returned to the colony room. Animals were closely monitored for health and recovery progress and received Meloxicam injections (20 mg/kg) to relieve pain for three additional days.

### Food restriction protocol

Before initiating behavioral training, mice were subjected to a food restriction protocol to increase motivation to perform the task. Briefly, mice were handled and weighed for at least two consecutive days before starting the food restriction protocol. Then, mice were food restricted to 2.5 g of chow per mouse per day and weighed daily to monitor maintenance to 85-90% of their predicted free-feeding weight based on average growth curve data for the strain (The Jackson Laboratory, Bar Harbor, ME). To familiarize the mice to Nesquik® strawberry milk (Nestlé, Vevey, Switzerland), which was used as reward during rCPT training, a 4×4 inch weighing plate (VWR, Radnor, PA, USA) containing ∼2 mL of strawberry milk was introduced to the home cage for three consecutive days. The weighing plate was left in the cage until all mice had sampled the strawberry milk.

### Behavioral training

#### Habituation

Mice were given two consecutive habituation sessions (20 min length) in Bussey-Saksida mouse touchscreen chambers (Lafayette Instruments, Lafayette, IN) to familiarize them to the chambers. In habituation sessions, 1 mL of strawberry milk was placed into the reward tray. The screen was responsive to touch, but touches were not rewarded.

#### Rodent Continuous Performance Test (rCPT)

rCPT training protocol was based on a previously described protocol [35]. Briefly, mice were trained in the touchscreen chambers, which were connected to a computer running ABET II software (Campden Instruments, Loughborough, UK) to track behavioral responses during rCPT sessions.

*Stage 1:* Mice received 45 min training sessions (Monday-Friday), during which they learned to respond to a visual stimulus (white square) presented at the center of the touch screen. The stimulus was displayed for 10 s (stimulus duration, SD), during which a touch in the center of the screen produced delivery of ∼20 uL of strawberry milk in the reward tray located on the opposite side of the chamber. Following SD, a 0.5 s limited hold (LH) period was given in which the screen was blank, but a touch would still yield a reward. Upon interacting with the stimulus, a 1 s tone (3 kHz) was delivered, the reward tray was illuminated signaling reward delivery, and the schedule was paused until a head entry into the reward tray was detected by an IR beam.

Then, a 2 s intertrial interval (ITI) would begin before the subsequent trial started. If the mouse did not interact with the stimulus during the SD or LH, an ITI would start, and the next trial would follow. The criterion for a mouse to advance to the next stage was to obtain at least 60 rewards per session in two consecutive sessions.

*Stage 2:* In Stage 2, a target stimulus (S+) was introduced. The S+ consisted of a square with either horizontal or vertical black and white bars that replaced the white square at the center of the screen. Sessions were 45 min long, and each mouse was assigned either horizontal or vertical oriented S+ for the remaining sessions of the experiment. The S+ assignment was counterbalanced. During Stage 2, the SD was reduced to 2 s, and LH was increased to 2.5 s. Interaction with S+ (hit) during SD + LH resulted in reward delivery. Once a mouse obtained at least 60 hits / session in two consecutive sessions, a recording session (see calcium endoscopic *in vivo* imaging) followed the day after (Stage 2 recording session). A mouse was moved to the next stage if it obtained at least 55 hits in the recording session. If a mouse failed to obtain at least 55 hits in the recording session, the recording session was repeated until the mouse obtained at least 55 hits.

*Stage 3:* In Stage 3, a non-target stimulus (S-) consisting of a snowflake shape presented at the center of the screen was introduced. On each trial, the probability of S+/S- was 50%/50%. The SD and LH were identical to Stage 2, but the ITI length was either 2 or 3 s in length (randomized ITI duration during trials). Similar to Stage 2, screen touches during S+ (hit) yielded a reward but not screen touches during S-(false alarm (FA)). A FA resulted in the beginning of the ITI followed by a correction trial. In correction trials, a S-was presented again. If another FA occurs, a new correction trial starts until the mouse doesn’t interact with the S-(correct-rejection). We used the discrimination index (d’), which is a measure of sensitivity bias ( perceptual discriminability between the S+ and S−) to determine attention performance during Stage 3. Mice were trained in Stage 3 until they reached a d’ score of 0.6 or higher for two consecutive days. A recording session occurred during the first Stage 3 session (Stage 3-early) and a second recording session occurred in the following session after two consecutive stage 3 sessions with a d’ of 0.6 or higher (Stage 3-late). In the case that during Stage 3-late a mouse had a d’ under 0.6, the recording session was repeated until they achieved a session with d’ of 0.6 or higher.

### Behavioral scoring

Behavioral databases containing the timestamps from stimuli presentation, hits, false alarms, latency to response, etc. were retrieved from ABET II (Lafayette Instruments, Lafayette, IN) and Whisker server (Cambridge University Technical Services, UK). Behavioral data was analyzed using Excel to obtain performance scoring parameters. Performance scoring parameters were similar to those described in [13] and [35]. Briefly, to assess attention performance during Stage 3 training, we calculated discrimination index **d’** with the following formula: d’ = z(hit rate) - z (FA rate)

Whereas:

hit rate (HR) = hits / hits + misses

FA rate (FAR) = false alarms / false alarms + correct rejections.

### Endoscopic in vivo calcium imaging

Single cell calcium transients were recorded though a GRIN lens coupled to a miniaturized microscope (nVista 3.0, Inscopix Inc. San Diego, CA) at three different behavioral training timepoints: 1) Stage 2; 2) Stage 3-early; and 3) Stage 3-late. Miniscope data was recorded at a 10 Hz frequency rate and the LED power, GAIN, and focus plane were optimized for each mouse. To identify calcium activity from individual neurons from raw miniscope recordings, videos were first preprocessed using Inscopix Data Processing Software (IDPS), Inscopix Inc. San Diego, CA). Videos containing calcium transients were spatially downsampled by a factor of 2, bandpass filtered (0.005 −0.5 pixels), motion corrected, and fluorescent values were normalized using a ΔF / F algorithm. Calcium transients for individual neurons were extracted using principal component analysis (PCA) using the following parameters: average diameter: 15-20 pixels, ICA convergence threshold: 0.00001, ICA temporal weights: 0, ICA max interactions: 100, Block size: 1000, ICA unimix dimension: Spatial. All identified neurons were visually inspected for soma-like morphology (size and shape) and all extracted traces were visually inspected for characteristic dynamics. Neurons with abnormal morphology or with non-consisting calcium transients were rejected for further analysis. Data was exported to the IDEAS platform (Inscopix Inc. San Diego, CA) and a built-in quality control algorithm was used to verify the quality of the extracted traces. Exclusion criteria for recordings were made based on quality control analysis on the ideas platform.

### Peri-event calcium imaging analysis

For the comparison of calcium activity surrounding behavioral responses (Hits and False Alarms [FA]), we used the Peri-Event-Analysis Workflow v4.3.0 tool in the IDEAS platform with the following parameters: visual window pre-event, 8 seconds; visual window post-event, 8 seconds; statistical window pre-start, 3 seconds; statistical window pre-end, 0 seconds; statistical window post-start, 0 seconds; statistical window post-end, 3 seconds; number of random shuffles, 1000; seed, 0; and significance threshold, 0.05. Each type of response was analyzed independently.

For each event-type, neural activity was first z-scored individually for each neuron. Δ activity was then computed by comparing the true peri-event activity to a null distribution generated by circularly permuting the event times relative to the neural time series. This permutation process was repeated 1000 times. For each neuron, a bootstrap probability was calculated as the fraction of the null distribution greater than the true Δ activity, and the fraction less than the true Δ activity. A two-tailed test was applied, comparing the resulting p-value to α/2 on each side of the distribution (α = 0.05). Neurons were classified as up-modulated if the bootstrap probability was less than α/2, down-modulated if greater than 1–α/2, and non-modulated if the bootstrap probability fell between α/2 and 1–α/2.

### Generalized Linear Models to assess recruitment across sessions and response type

Generalized Linear Mixed Models (GLMMs) were employed to examine the recruitment of modulated neurons across different sessions, modulations, and event alignments. All analyses were conducted in MATLAB using the fitglme function. The GLMMs were used to assess how proportions of modulated neurons (Up, Down, and Non) varied across behavioral sessions (Stage-2, Stage 3-early, and Stage 3-late), response type (Hit and FA), and event alignments (stimulus presentation versus screen touch). In all models, the dependent variable was the ResponseCount, which was calculated as the product of the proportion of neurons and the total number of neurons recorded in each session. The GLMMs incorporated Session, Modulation, and Alignment (stimulus or screen touch) as fixed effects and included random intercepts for Mouse to account for individual differences in neuronal recruitment across animals. For each model, the Gaussian distribution was used to fit the data with an identity link function. The models included interaction terms between session, modulation, and alignment to assess how the effects of modulation and alignment varied across different sessions. The fit quality was assessed through model fit statistics, including the Akaike Information Criterion (AIC) and Bayesian Information Criterion (BIC). For hypothesis testing, we used ANOVA to examine the significance of the main effects and interactions, with post-hoc comparisons to further assess differences between specific conditions. Statistical significance was evaluated with p-values < 0.05, and coefficients were reported with 95% confidence intervals. Code is available upon request.

### Detecting neural states using Hidden Markov Models

HMMs have been previously validated for detecting neuronal states associated with behavioral outcome [36]. All analyses were performed in Python 3 using custom scripts. Calcium-imaging traces for each animal were truncated to the first 45 minutes of recording and traces were standardized (z-scored) to zero mean and unit variance, using the *zscore* function from the *scipy.stats* package. To reduce high-frequency noise without introducing lag, a 1-second moving-average filter (10-frame window, center-aligned) to the z-scored data was applied using the *rolling* function from the *pandas* package. Dimensionality reduction was then performed on the z-scored, smoothed data via principal component analysis (PCA), using the *pca* function from the *sklearn.decomposition* package. The first 45 principal components which together explained the majority of variance were retained, and recorded the cumulative variance explained as a quality check. A two-state (k=2) Hidden Markov Model (HMM) was applied to the first 10 PCs of each animal using a Gaussian HMM with diagonal covariance [37–39], using the *hmm* function from the *hmmlearn* package. For model fitting, an EM algorithm up to 200 iterations (default tolerance) was run.

### Hidden Markov Model Assessment and Validation

The Hidden Markov Model (HMM) requires specification of the hyperparameter *K*, which defines the number of states the model will resolve from the data. To validate our selection of *K* = 2 (i.e., a two-state model), we computed the log-likelihood for models with one, two, and three states. Log-likelihood values were obtained using the built-in model.score function from the *hmmlearn* package.

To determine model performance, accuracy, and assess potential overfitting, we performed three-fold cross-validation on the two-state hmm model for each animal and each session of S3 Good performance. Three-fold C.V. involved splitting each session into four equal parts, and defining test/train data depending on the iteration of C.V. For one-fold C.V. the training set is the first 25% of the data and the test set is the second 25%. For two-fold C.V. the training set is the first 50% of the data and the test set is the third quarter of the data, etc. To compare accuracy, the model was trained and fitted on the training set, and those same parameters were applied to fitting on the test set, upon which the log-likelihood was computed for both sets and compared. We applied cross-validation in this continuous manner to avoid skewed model accuracy results stemming from disruption potential network dynamics evolving across the session (i.e. model inaccuracy from training on future data and testing on previous data).

### Posterior probabilities

For determining posterior probabilities of responses happening in neuronal states, behavioral timestamps for Hits, Mistakes, and their logical OR (“Any”) were converted to 10 Hz frame indices and binarized into event vectors E(t)∈{0,1}. Given the HMM-decoded state sequence state(t) ∈ {0,1}, we then computed—for each event type *E* and each state i ∈ {0,1}—P(State_i_ | Hit) = Σ_t: Hit_ 1[state(t) = i]) / total Hits i.e. the fraction of all event-frames that occurred while in *state i*. These six posterior probabilities, one for each combination of state and event type, were tabulated per animal.

### Calcium activity correlation analysis during states

To determine the calcium activity correlations during behavioral states, time lapses belonging to each state were input into the *Neural Circuits Correlations v6.0.2-6.0.1* tool in the *IDEAS platform*. For each mouse, the Neural Circuits Correlations v6.0.2-6.0.1 too collapses all frames from each state and then computes pairwise Pearson correlation between all neuron pairs in each state and creates an n×n correlation matrix (where n = number of neurons) using standard Pearson correlation. Pearson correlation coefficients are calculated as: r = Σ[(Xi - X^-^)(Yi - Ȳ)] / √[Σ(Xi - X^-^)² × Σ(Yi - Ȳ)²], where Xi and Yi are individual trace values, and X^-^ and Ȳ are the means. Self-correlations (diagonal values) values are adjusted to zero and correlation matrices are sorted by grouping similar cells together while preserving the identity of the neurons. Spatial information regarding the Euclidean distance between pairs is calculated and then used for determining decay of correlated activity between pairs in each state using custom MATLAB algorithms. Additionally, for each cell, the Neural Circuits Correlations v6.0.2-6.0.1 tool computes the maximum correlation value across its positive and negative (anticorrelation value) values with other cells. Cell-cell correlation data across mice is then combined and analyzed using the *Compare Neural Circuit Correlations Across States* tool in the Ideas platform that compares average positive/negative correlations across states using one-way Repeated Measures ANOVA. For cell level analysis across states, the tool uses Linear Mixed Models (LMM) to compare the cell-level statistics across states, accounting for within-subject variability and potentially group differences, handling nested structure (cells within subjects). Additionally, pairwise comparisons following significant ANOVA or LMM results were performed to pinpoint differences between specific states or groups. Cumulative probability plots were created with maximum correlation values (positive and negative).

### Analysis of correlation decay as a function of distance

To investigate whether spatial decay of correlated neuronal activity differs between cognitive states, we computed pairwise Pearson correlations of calcium activity across neurons during the Engaged and Disengaged states. Correlation values were categorized as positive, negative, or non-significant based on thresholds obtained via a permutation-based null distribution, using 10,000 random sign-flip iterations. For each category and state, we then fit an exponential decay model of the form *y = A·exp(k·x)* to examine how correlation strength declines with inter-neuronal distance. Where *y* is the observed correlation strength between neuron pairs, *x* is the physical distance between those neurons, *A* is the initial correlation strength at zero distance, and *k* is the decay rate constant (with more negative values indicating faster spatial decay). To statistically assess differences in decay rate (*k*) between states, we derived 95% confidence intervals for each fit using bootstrapping and conducted z-tests for independent exponential parameters.

## Results

### Experimental design and acquisition of task proficiency in the rCPT during in vivo calcium imaging sessions

Changes in PrL neuronal activity are linked to sustained attention [30,34,40]. However, the cellular mechanisms in the PrL that contribute to sustained attention are not fully understood. To investigate neuronal activity patterns during sustained attention, we expressed a viral vector encoding GCaMP6f in PrL neurons, and then trained mice on the rCPT. We recorded single cell calcium activity in PrL neurons during three rCPT sessions that differ in cognitive demand and task proficiency: 1) Stage 2: mice are proficient in detecting a visual stimulus (low cognitive demand), 2) Stage 3-early: mice are required to discriminate between two stimuli, but performance is close to chance (high cognitive demand/low proficiency), and 3) Stage 3-late: mice are required to discriminate between stimuli, and performance demonstrates the ability to discriminate between the S+ and S-(high cognitive demand/high proficiency) (Fig. 1A-B). Mice quickly learned to respond to the stimulus presented during Stage 1 (Fig. 1C), and displayed a high number of correct responses since the first Stage 2 session (Stage 2 first session hits: 105. ± 14.01, Stage 2 last session hits: 87.08 ± 7.03; Fig. 1D). Mice reached criteria for Stage 2 between the second and fourth Stage 2 session. A decrease in the number of responses was observed in some mice following tethering, but these mice resumed criteria for performance in the following recording session (Supplementary Fig. 1). As expected, in the first Stage 3 session (Stage 3-early), mice exhibit low task performance (low d’ score, and similar number of correct responses (hits), and incorrect responses/False Alarms (FAs); Fig. 1E-J), but quickly improved, reaching criteria between the 6th and 10th session of Stage 3-late (Stage 3-early d’= −0.02 ± 0.07 vs Stage 3-late d’= 0.88 ± 0.05, Fig. 1E and H. Stage 3-early hits= 32.92 ± 5.63 vs Stage 3 late hits= 59.50 ± 8.0, Fig. 1F and I. Stage 3-early FAs= 32.25 ± 4.1 vs Stage 3 late FAs= 17 ± 3.27, Fig. 1G and J). Notably, response latency after stimulus presentation did not significantly differ between hits and FAs in Stage 3-early (Hits: 1.82 ± 0.08 s vs FA 1.82 ± 0.08 s; Fig. 1K), however during stage 3-late, we identified a reduction in the latency of FAs compared to hits (Hits: 1.8 ± 0.08 s vs FA 1.6 ± 0.08 s; Fig. 1K).

**Figure 1.**
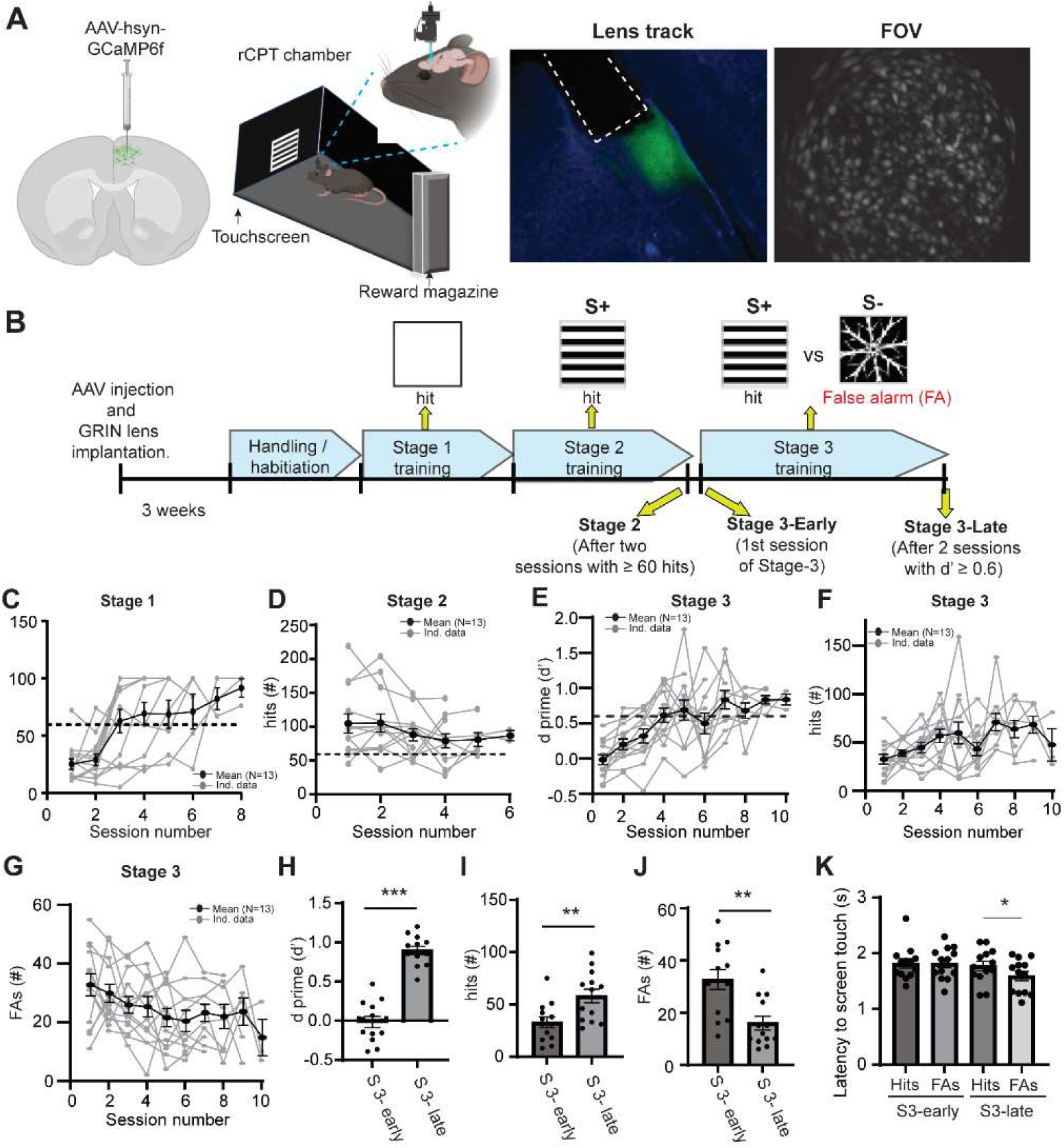
Behavioral performance during rCPT training sessions. A) Schematic of experimental strategy for *in vivo* calcium imaging of PrL neurons during rCPT. B) Timeline for calcium imaging sessions during rCPT behavioral training. C) Number of correct responses (hits) during Stage 1. D) Number of hits during Stage 2. Blue dots represent calcium imaging sessions during Stage 2. E) Time course of behavioral performance (represented as d’) during Stage 3 sessions. F) Time course of the number of hits during Stage 3. G) Time course of the number of false alarms (FAs, incorrect responses) per session during Stage 3. H) Bar plot comparing performance between Stage 3 (S3)-early and Stage 3 (S3)-late. Mice showed a significant increase in d prime values during S3-late, paired t test: t_12_ = 12.62 p˂ 0.0001 I) Bar plots showing the number of hits during S3-early and S3-late. There is a significant increase in the number of hits during S3-late, paired t test t_12_= 3.60 p= 0.0042 J) Number of FAs during S3-early and S3-late. There is a significant decrease in the number of hits during S3-late, paired t test t_12_= 1.066 p= 0.32). K) Bar plot showing latency to hits and FA during S3-early and S3-late. The latency to respond during FA in S3-late is significant lower than to hits, paired t-test t_12_= 2.75 p= 0.017

### Increased calcium activity in PrL neurons during correct responses

To determine the neuronal dynamics of PrL neurons during rCPT, we first analyzed peri-event calcium activity in time windows surrounding hits or FAs across different rCPT sessions aligning the events to behavioral responses (screen touches). In Stage 2, 18.7% (341/1822) of PrL neurons were up-modulated, 23.4% (426/1822) were down-modulated, and 57.9% (1055/1822) were non-modulated (Fig. 2A-C). In Stage 3-early, 15.2% (282/1860) of neurons were up-modulated, 20.7% (385/1860) were down-modulated, and 64.1% (1193/1860) were non-modulated during hits (Fig. 2A-C). In contrast, during FAs, only 10.5% (196/1860) of neurons were up-modulated, 10.4% (194/1860) were down-modulated, and 79% (1470/1860) were non-modulated (Fig. 2A-C). Finally, during Stage 3-late, 17.4% (375/2146) of PrL neurons were up-modulated, 31.4% (675/2146) were down-modulated, and 51.1% (1096/2146) were non-modulated during hits. During Stage 3-late FAs, 6.4% (138/2146) of PrL neurons were up-modulated, 7.7% (165/2146) were down-modulated, and 85.9% (1843/2146) were non-modulated (Fig. 2A-C). It is important to highlight that most of the modulated neurons in Stage 3-early and Stage 3-late were specific to behavioral response type (hits or FAs), with only a minority of neurons modulating their activity during both hits and FAs (Fig. 2D).

**Figure 2.**
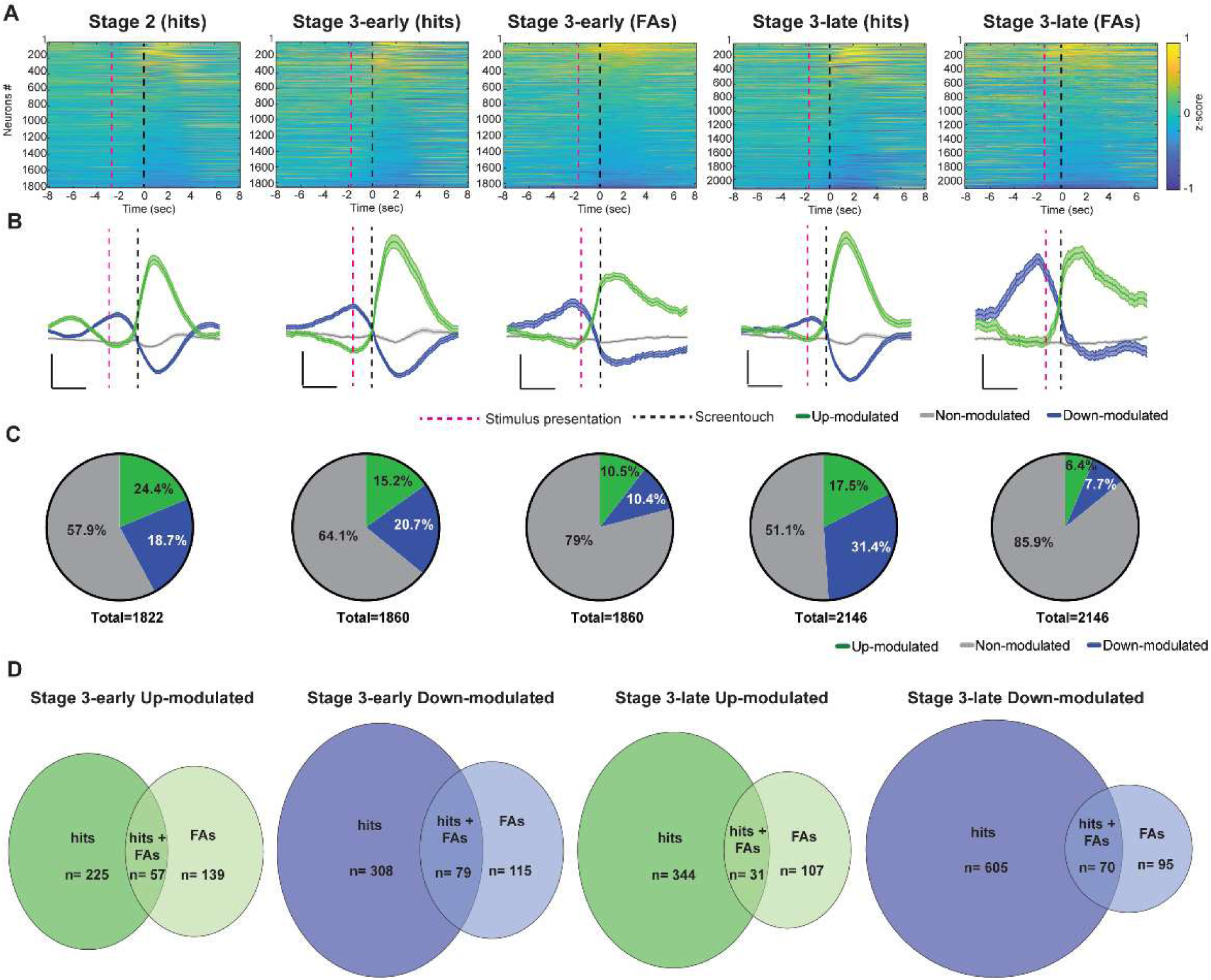
Calcium activity increases in PrL neurons during correct responses in proficient sessions. A) Normalized change in calcium activity (z-scored) surrounding hits (correct responses) and false alarms (FAs, incorrect responses) during rCPT recording sessions. B) Calcium activity traces from up-modulated (green), down-modulated (blue) or non-modulated (gray) neurons surrounding the behavioral response. The red dotted line represents the stimulus presentation and the black dotted line represents the behavioral response (screen touch). Scale bar: 2 s, and z-score= 0.2. C) Pie charts showing the proportion of neurons that were up-modulated (green), down-modulated (blue) or non-modulated (gray) surrounding the behavioral response. D) Venn diagrams showing up-modulated (green) and down-modulated (blue) neurons by hits, FAs, and hits and FAs across Stage 3 recording sessions.

While analyzing calcium activity of individual neurons during behavioral responses, we noted that calcium activity began to increase between stimulus presentation and the screen touch in a subset of up-modulated neurons (Fig. 2A). This increase in calcium activity is reflected as a change in the slope of the averaged calcium transient prior to screen touch (Fig. 2B). Additionally, the average calcium transient in down-modulated neurons across sessions displayed a biphasic pattern: calcium activity increased surrounding stimulus presentation (positive slope), followed by a decrease after screen touch (negative slope) (Fig. 2A and B). This negative slope after screen touch appeared blunted during FAs (Fig. 2B). Since we observed changes in calcium activity preceding screen touch, we reanalyzed calcium activity during hits and FAs, but aligned the events to the stimulus presentation. We found similar response patterns to those observed when centering the analysis to the screen touch (Supplementary Fig. 2). A GLM model revealed no significant differences in the proportion of modulated neurons across responses when the alignment of the events was changed (ANOVA F-stat_1,37_= 2.3496; p= 0.126).

To compare whether recruitment of up- or down-modulated neurons varied significantly across sessions, we ran a GLM model comparing the change in the proportion of modulated versus non-modulated neurons during Hits across Stage 2, Stage 3-early, and Stage 3-late sessions. We found a significant increase in the proportion of modulated neurons during Stage 3-late compared to Stage 3-early (F-stat_2,74_ = 12.927; p ˂ 0.001). However, no significant difference was observed when comparing recruitment between Stage 2 and Stage 3-late (Z-value = 0.58448, p= 0.5589). Additionally, we tested whether recruitment changed significantly based on response type (Hit vs FA) during Stage 3 sessions (Stage 3-early and Stage 3-late). We found significant effects of stimulus type and modulation status, with neurons being more likely to be modulated during hits than during FAs (F-stat_1, 97_ =19.594 p˂ 0.001). However, we did not find significant differences in recruitment during FA between Stage 3-early and Stage3-late (F-stat_1, 97_= 0.163 p= 0.687). Together, these findings demonstrate that neuronal recruitment during the rCPT is higher during hits, with an increase in recruitment associated with task proficiency but not cognitive demand.

### Responsive and non-responsive periods of task engagement are intercalated during the rCPT

A key characteristic of sustained attention is its endurance over time. In humans, sustained attention fluctuates across time resulting in attentional lapses [41]. We hypothesized that attention lapses in the rCPT could be reflected by alterations in task engagement within sessions. To assess changes in task engagement during rCPT behavior, we analyzed response patterns by plotting behavioral responses (hits and FAs) over time within sessions. This qualitative analysis revealed a heterogeneous distribution of responses, with responsive periods interspersed with non-responsive periods of variable length, characteristic of individual mice (Fig. 3A and Supplementary Fig. 3 and 4). Non-responsive periods were clearly observed as peaks when plotting the latency between behavioral responses (i.e., time elapsed between hits and FAs), and they were present across all sessions (Stage 2, Stage 3-early, and Stage 3-late) (Fig. 3B and Supplementary Fig. 4). We hypothesized that responsive and non-responsive periods reflect variations in task engagement, and that neuronal activity within the PrL reflects these changes. We reasoned that using unbiased methods to categorize global PrL neuronal activity within a session might identify periods where task engagement differs during the rCPT. To examine changes in global PrL activity associated with task engagement, we ran a Hidden Markov Model (HMM) with two states, hypothesizing that one state would be preferentially associated with responsive periods or engagement, while the other would correspond to non-responsive periods or disengagement. To validate the model, we compared the performance of the two-state HMM against models with one and three states by calculating the log-likelihood. Both the two- and three-state models outperformed the one-state model, indicating that they better captured the data structure. However, there was no significant difference in performance between the two- and three-state models (Supplementary Fig. 5). We next calculated the probability of behavioral responses occurring within each identified neuronal state for individual animals.

**Figure 3.**
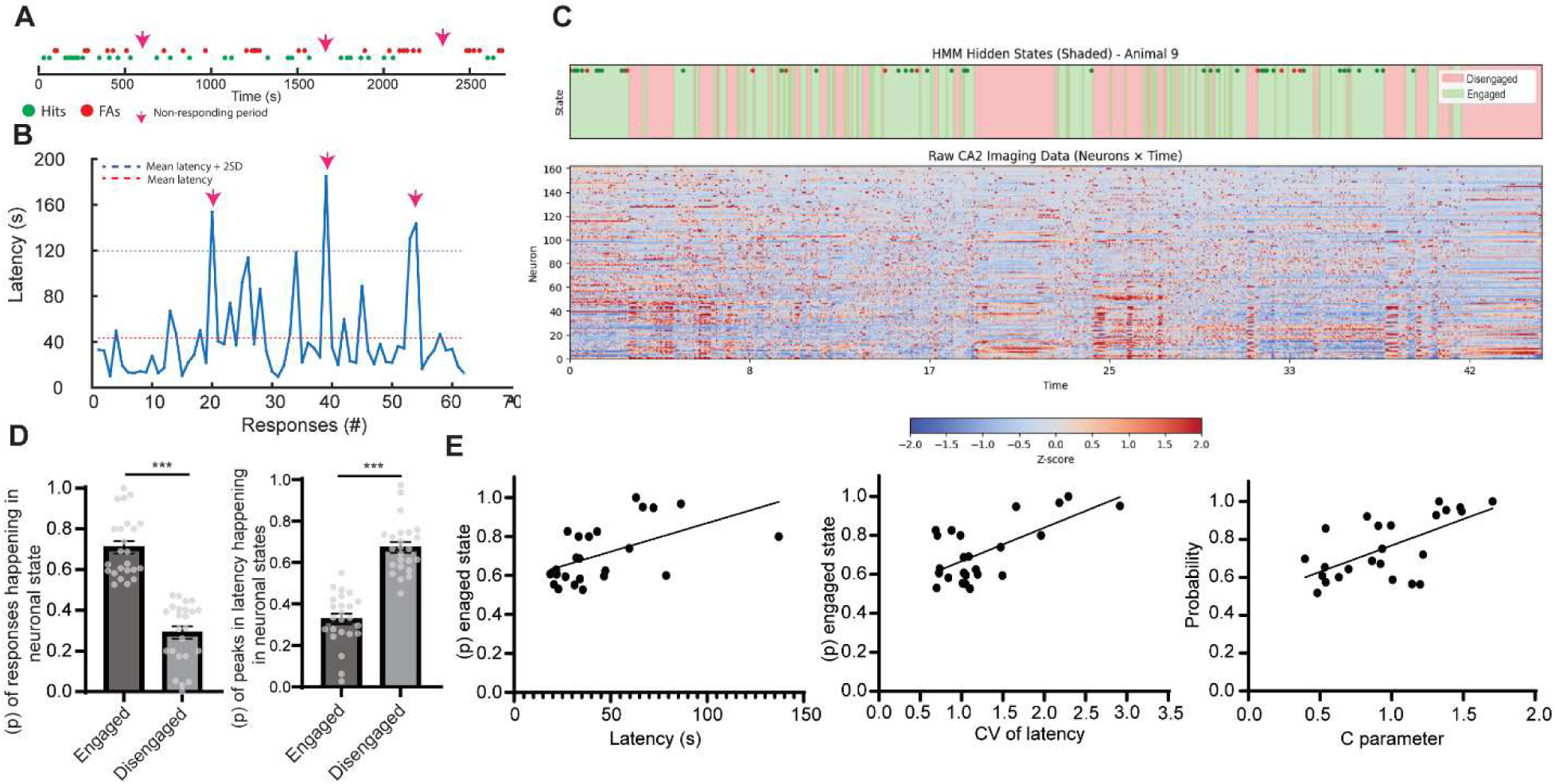
Neuronal states associated with task engagement. A) Timelines of a rCPT session showing heterogeneous pattern of responses (green dots represents hits while red dots represents FAs) across time. Arrows point to periods with no behavioral responses. B) Analysis of latency between responses during a rCPT session. Latency between responses is calculated as the time between two active responses (hits or FAs). Pink arrows point to peaks of latency between responses corresponding to the same periods indicated in A. (Red dotted line: mean latency between responses for that particular session. Blue dotted line; mean of latency +2 SD). C) Representative HMM analysis with two latent states on global PrL activity during a rCPT session. Periods corresponding to engaged (green) and disengaged (pink) states aligned with behavioral events (hits= green circles, FAs= red circles) (top) and time course of global PrL activity during rCPT session (Bottom). D) Bar graphs showing the probability of responses happening in each latent state (left) and the probability of the time periods corresponding to peaks in latency between responses happening in each state (right). The probability of responses happening in the engaged state is significantly higher to the disengaged state (t_46_= 9.64 p˂ 0.001) while the probability of the time periods corresponding to peaks in latency between responses happening in the disengaged state is higher (t_46_= 9.45 p˂ 0.001. E) Correlation graphs showing probability of responses happening in the engaged state over the mean latency between responses (left) (pearson r= 0.535, p= 0.007), coefficient of variation (CV) for the mean latency between responses (middle) (pearson r= 0.657, p˂ 0.001), and the *c* parameter (right) (pearson r= −0.6343, p˂ 0.001).

Across mice, responses were consistently more likely to occur in one particular state, which was labeled as the engaged state (engaged state: 0.71 ± 0.03; disengaged state: 0.29 ± 0.03) (Fig. 3C–D). Similarly, when we assessed the probability of time periods corresponding to peaks in response latency—previously identified as non-responsive periods (Supplementary Fig. 4)—we found that these periods occurred significantly more often in the other state, which we labeled as disengaged state (engaged state: 0.33 ± 0.03; disengaged state: 0.67 ± 0.03) (Fig. 3C–D), supporting the association of neuronal states with task engagement. Interestingly, the probability of responses occurring in the engaged state was significantly higher in the two-state compared to the three-state HMM (Supplementary figure 6).

Although the probability of responses happening in the neuronal state links them with task engagement, this probability varied across mice without a significant effect between session type (Stage 2, Stage 3-early, and Stage 3-late; Supplementary Fig. 7). We hypothesized this probability might reflect how well the HMM identified states related to task engagement. If this hypothesis is correct, we would expect the probability of responses happening in the engaged state to be higher in mice with clear changes in task engagement — characterized by periods with high variability in latency between responses (reflecting intercalated periods of low and high latency) elevated values of the *c* parameter, which reflects response strategy. Notably, higher *c* values could indicate the presence of attentional lapses during the rCPT session. Alternatively, the HMM might differentiate states driven by the recruitment of neurons during behavioral responses (i.e., hit and FA responses may recruit a significant number of neurons, aiding in state identification). In this case, we would expect that the probability of responses happening in the engaged state correlates with the number of responses or response type. To test these hypotheses, we analyzed the correlation between the probability of responses happening in the engaged state with the latency, the coefficient of variation of latency, the *c* parameter, and the number and type of responses across mice. We found that the probability of responses happening in the engaged state correlated with the latency between responses (Pearson r= 0.535, p= 0.007), the variability in that latency (coefficient of variation) (Pearson r= 0.657, p˂ 0.001), and the *c* parameter (Pearson r= 0.634, p˂ 0.001) (Fig. 3E), but not with the total number of responses (Pearson r= −0.207, p= 0.33) or response types (Pearson r (hit)= −0.144 p= 0.502; Pearson r (FA)= −0.194, r= 0.363) (Supplementary Fig. 8). These findings suggest that when a mouse shows clear differences in task engagement as reflected by high variability in the latency between responses due to alternating responsive and non-responsive periods, the model is better able to identify neuronal states associated with task engagement using PrL neuronal activity.

### Network changes in PrL calcium activity during neuronal states associated with task engagement

Following HMM analysis of PrL activity and the identification of neuronal states associated with task engagement, we sought to characterize differences in network-level activity between these states. To this end, we analyzed correlations in calcium activity between PrL neurons during the engaged and disengaged states. Across mice, we observed distinct correlation profiles in these states (Fig. 4A), with an overall reduction in PrL calcium activity correlations during the engaged state. Specifically, cumulative probability curves for positively and negatively correlated values, as well as for the maximum correlated value per neuron, were shifted to the left in the engaged state (Fig. 4B–D). Notably, we found that all types of correlations between neuron pairs declined as the distance between pairs increased (Spearman’s ρ for positive correlations—engaged: −0.06, *p* < 0.001; disengaged: −0.06, *p* < 0.001; negative correlations—engaged: −0.12, *p* < 0.001; disengaged: −0.10, *p* < 0.001; Supplementary Fig. 9).

**Figure 4.**
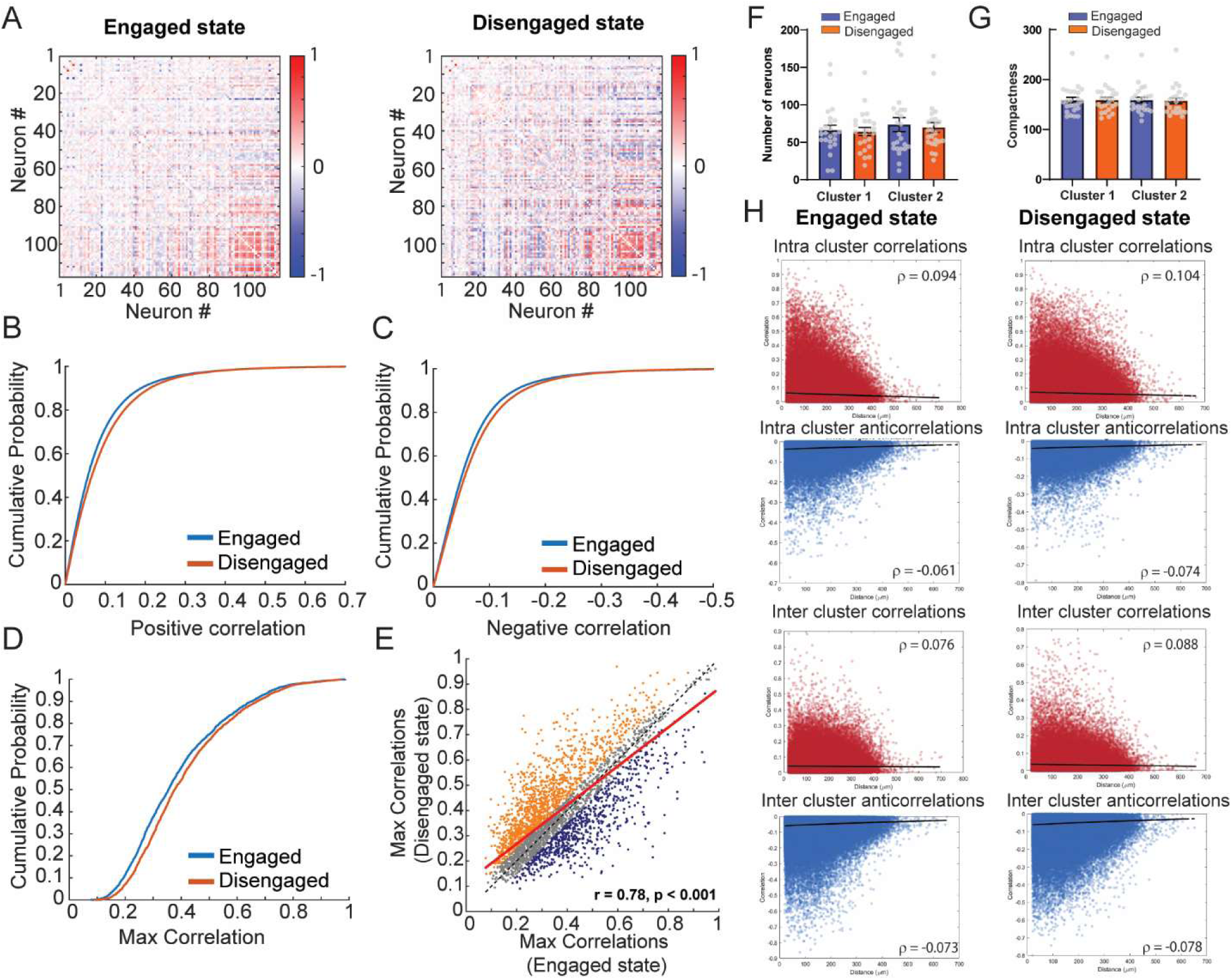
Characterization of PrL calcium activity during neuronal states associated with task engagement. A) Correlation matrices of neuronal activity from a representative rCPT session. B) Cumulative probability plots showing positive correlation values of neuronal activity across engaged and disengaged states during rCPT (KS test D = 0.0491, p ˂ 0.001, n_engaged_ = 163,755 pairs, n_disengaged_ = 162,166 pairs) C) Cumulative probability plots showing negative correlation (anticorrelations) values of neuronal activity across engaged and disengaged states during rCPT (KS test D = 0.0630, p ˂ 0.001, n_engaged_ = 16,2089 pairs, n_disengagedt_ = 163,678 pairs). D). Cumulative probability plots showing maximum correlation values in neuronal activity across engaged and disengaged states during rCPT (KS D = 0.0884, p ˂ 0.001, n_engaged_ = 3643, n_disengaged_ = 3643). E) Analysis of the maximum correlation value in engaged vs disengaged states. Neurons in orange exhibit higher correlation during non-dominant state compared with dominant state. F) Bar graph showing number of neurons belonging to identified clusters across neuronal states (Cluster 1: engaged= 66.2 ± 6.6 neurons, disengaged = 64.3 ± 5.5 neurons; Cluster 2: engaged= 73.6 ± 9.3 neurons, disengaged = 69.8 ± 6.5 neurons). G) Bar graph showing spatial compactness of the identified clusters across neuronal states (Cluster 1: engaged= 158.7 ± 5.6, disengaged = 159.1 ± 5.3; Cluster 2: engaged= 159.9 ± 5.5, disengaged = 157.5 ± 5.9). H) Analysis of neuronal activity correlation between pairs of neurons and distance show state dependent decrease in correlated activity with physical distance between pairs of neurons.

To further investigate how activity correlations change across states, we analyzed the maximum correlated values of individual neurons and identified those neurons having preferentially correlated activity in one state over the other (Fig. 4E), suggesting the existence of cell-pair correlations linked to specific network states. To assess whether correlated activity patterns reflect distinct network organization, we performed K-means clustering on the correlation matrices for each state. The optimal number of clusters (K) was determined using silhouette score analysis, which indicated that *K* = 2 was most appropriate in the majority of cases (Supplementary Fig. 10). After identifying the clusters in each state, we examined changes in their neuronal composition and spatial distribution. We found no significant differences in cluster sizes (F_3,92_= 0.336 p=0.799) or spatial compactness F_3,92_= 0.0185 p=0.996) across states (Fig. 4F and G). However, when we analyzed the decay of correlated activity with physical distance between neurons—both within (intra-cluster) and between (inter-cluster) clusters—as a measure of how behavioral states affect functional coupling within and across cell-pair correlations, we observed significant state-dependent changes. Specifically, intra-cluster positive correlations decreased in the engaged state (paired *t*-test, *t*_₈₂,₈₀₉_ = −20.04, *p* < 0.001), while intra-cluster negative correlations became more pronounced (*t*_₆₂,₀₇₆_ = 18.23, *p* < 0.001). Similarly, inter-cluster positive and negative correlations were both significantly higher in the disengaged state (inter-cluster positive correlations: *t*_₇₂,₆₀₈_ = −26.22, *p* < 0.001; inter-cluster negative correlations: *t*_₉₆,₁₄₆_ = 16.11, *p* < 0.001) (Fig. 4H). Interestingly, the steeper decay in correlations is also present when significant positive and non-significant correlations between pairs of neurons were analyzed separately (Supplementary Fig. 11). These findings suggest a state-dependent reconfiguration of functional connectivity within the PrL network that tracks changes in task engagement.

## Discussion

In this study, we used *in vivo* calcium imaging to characterize activity patterns of PrL neurons during sustained attention in the rCPT. Our results revealed heterogeneous responses of PrL neurons across behavior, with a higher proportion of PrL neurons modulated during correct responses, especially in proficient sessions (Stage 2 and Stage 3-late). These results are in line with previous electrophysiological studies using single unit recordings in fully trained mice that showed a higher proportion of modulated PrL neurons during trials that ended in correct responses [29,31,40]. Like other attention tasks, the rCPT assesses attention by determining the animal’s ability to discriminate between stimuli over extended sessions where at least one stimulus is paired with reward [13]. In addition to sustained attention [17,29,30,33,42], the PrL is linked to stimulus discrimination [43–45] and reward processing [46,47,48], which are important for rCPT performance. In our data, aligning the events to behavioral responses (screen touch) or stimulus presentation show similar response patterns in the calcium activity, where the peak of PrL calcium transients occurred between the stimulus presentation and directly following screen-touch, possibly reflecting PrL computations for visual discrimination, decision making, and reward processing, all of which are consistent with a role for the PrL in attention-related sensory processing [43,49,50].

Unlike previous studies that assessed neuronal patterns in attention tasks in fully trained animals [29–31,33,42,51], we analyzed PrL calcium dynamics across different stages of rCPT training, encompassing varying levels of cognitive demand and task proficiency. We observed response-specific changes in PrL calcium activity in proficient and non-proficient sessions, consistent with our previous findings of increased PrL delta and theta power during correct responses across training stages [34]. Together, these data suggest that calcium activity in PrL neurons and LFPs in the PrL track stimulus discrimination, decision making, and response outcome across learning states during periods of certainty and uncertainty [52–54].

Although the proportion of PrL neurons that were modulated during incorrect responses was small, we identified a subset whose calcium activity peaked close to the time of stimulus presentation and others that peaked near after screen touch. It is possible that neurons with peak calcium activity close to the time of stimulus presentation may participate in cognitive processes related to behavioral control during incorrect responses such as response inhibition. Previous reports have shown a role of the PrL in conflict monitoring [55] and response inhibition, as lesioning [15,56] or inactivating the PrL [57] impairs accuracy, and results in a disinhibited response profile on attention tasks. Other possibilities are that these neurons represent preparatory activity such as reward expectation or reflecting simple habitual responses emerged from training, functions which have been previously related to the PrL [58,59]. For neurons with peak activity after the screen touch, it is possible that they participate in outcome monitoring and their activity increased as a result of an unexpected outcome (reward not being delivered). Previous evidence pointed to the PrL to be involved in encoding **r**eward prediction errors when unexpected negative outcomes occur [60–62]. However this idea contrasts with a previous report that showed that in rats, the ACC and not the mPFC exhibit phasic changes following behavioral responses [29]. This divergence may reflect species-specific differences (rat PFC vs. mouse PrL), but could also arise from differences in task structure and recording methodology. Totah et al. used single-unit recordings during a three-choice serial reaction time task with long inter-trial intervals, allowing for clear separation of neuronal activity phases. In contrast, the lower temporal resolution of calcium imaging, combined with shorter ITIs and rapid response latencies in our task, limits the ability to disentangle preparatory from post-response activity. Future experiments designed to temporally dissociate response components will be essential to clarify the precise role of mouse PrL neurons during incorrect responses.

### PrL activity patterns during attention lapses

Although peri-event analysis during correct and incorrect responses has provided important information about the role of specific brain regions and their activity patterns during processes related to sustained attention [29],[34,63], analyzing moment-to-moment fluctuations in global neuronal activity across entire sessions may offer a complementary approach to uncover the cellular mechanisms underlying variations in task engagement. In this study, we applied a Hidden Markov Model (HMM) to global PrL calcium activity as an unbiased computational method to identify latent neuronal states potentially linked to task engagement. HMMs infer discrete hidden states that best explain temporal patterns in population activity and enable the estimation of moment-to-moment state transitions, capturing the dynamic structure of neural activity across sessions.

Notably, HMMs have been used to infer latent neural states and have proven effective in detecting population dynamics related to behavioral processes, including decision-making, motor planning, and internal states [37–39,64]. To our knowledge, this is the first time that PrL global neuronal activity patterns analyzed through a HMM are used to associate task engagement during rCPT performance, which provided a significant advantage over supervised behavioral classification of task engagement that rely on event alignment to particular behavioral events such as omissions [65–69] or arbitrary thresholds. Importantly, while HMM provides a powerful, data-driven framework, several critical considerations must be acknowledged. First, the number of states inferred by the model is a hyperparameter that must be set a priori. Based on our data and hypotheses, we selected a two-state model, but it is plausible that additional states exist and may reflect other behaviorally relevant processes. Indeed, we observed inter-animal variability in the likelihood that the engaged state aligned with behavioral responses. While this variability may reflect individual differences in attentional engagement, it may also suggest that additional, unmodeled neuronal states contribute to task performance, thereby limiting the interpretive power of a two-state model. HMM state transition inferences across time are discrete rather than continuous, while this improves interpretability, it is to some degree an oversimplification as neural networks evolve in a more continuous manner [70]. Future directions may be to apply more flexible models such as dynamical systems-based modeling to capture these temporal dynamics. However, these more nuanced models also harbor reductions in interpretability.

While further refinements in the application of HMMs could enhance the study of neuronal states associated with task engagement, our characterization of the engaged and disengaged states revealed important insights. We observed distinct patterns of correlated activity between states, demonstrating population-level changes in the degree of PrL neuronal coordination that varied with engagement. Importantly, we do not interpret these state changes as reflecting just physical disengagement from the task, as our previous work demonstrated that mice remain oriented toward the stimulus on the vast majority of trials, even during periods of reduced performance [71]. Changes in correlated activity between states is in line with other data showing that attention causes cortical desynchronization by decreasing inter-neuronal correlation [72,73]. Assessing the role of different neurotransmitter systems in driving the changes in neuronal states correlated activity across different moments of task engagement could help to determine the cellular and molecular mechanisms underlying sustained attention. Norepinephrine (NE) signaling within the PrL-LC circuit is particularly interesting in this context. The PrL sends projections to the LC, which modulate noradrenergic transmission [74,75]. The PrL also receives noradrenergic inputs from the LC, which have a modulatory effect on attention [76,77,76–78]. NE released in the cortex facilitates synchronization during attention-related behaviors [79,80], and denervating noradrenergic inputs to the PrL decreases arousal and wakefulness in novel environments where attentional resources should be allocated [81]. Moreover, changes in LC-NE neuron activity are associated with changes in task engagement [82] as well as attentional lapses [80]. Better understanding on the regulation of bidirectional activity in the PrL-LC circuit, and the effect of NE release on neuronal activity in PrL neurons during changes in task engagement could advance our understanding on mechanisms underlying sustained attention.

---

**Supplementary Fig. 1.**
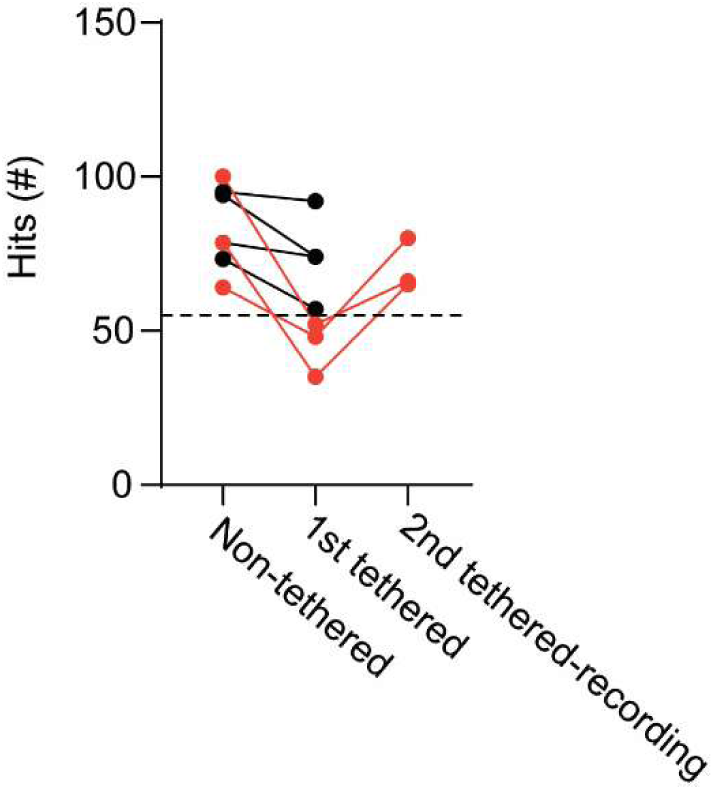
Effect of tethering on behavior during Stage 2 training sessions. Number of responses during a non-tethered session and tethered sessions. Mice represented by orange circles decreased their responses under criteria threshold (55 responses) during the first tethered session, and required an extra tethered session to reach criteria for recording.

**Supplementary Fig. 2.**
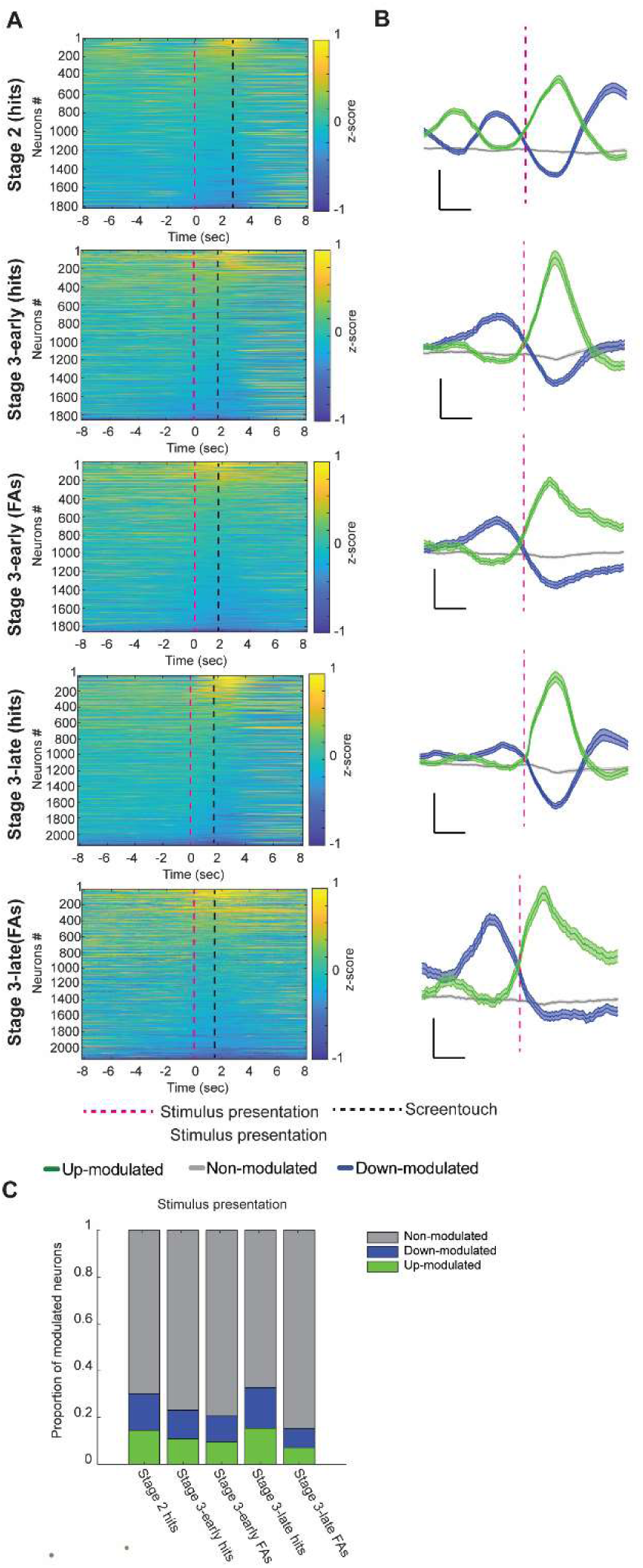
Peri-event analysis of PrL calcium activity during rCPT aligning events to stimulus presentation. A) Heat maps showing normalized change in calcium activity (z-scored) surrounding hits (correct responses) and false alarms (FAs, incorrect responses) during rCPT recording sessions after aligning events to stimulus presentation. B) Calcium activity traces from up-modulated (green), down-modulated (blue) or non-modulated (gray) neurons surrounding the behavioral response. The pink dotted line represents the stimulus presentation Scale bar: 2 s, and z-score= 0.2. C) Bar graphs showing proportion of neurons that were up-modulated (green), down-modulated (blue) or non-modulated (gray) surrounding the behavioral response. Stage 2 hits up-modulated = 14.5% (265/1822), down-modulated = 15.53 (283/1822), non-modulated = 69.92% (1274/1822); Stage 3-early hits up-modulated = 11.0% (204/1860), down-modulated = 12.2% (227/1860), non-modulated = 76.83% (1429/1860; Stage 3-early FAs upmodulated = 9.7% (180/1860), down-modulated = 11.1% (206/1860), and non-modulated = 79.3% (1474/1860); Stage 3-late hits upmodulated = 15.4% (331/2146), down-modulated = 17.4% (373/2146), non-modulated = 67.2% (1442/2146); Stage 3-late FAs up-modulated = 7.2% (154/2146), down-modulated = 8.1% (173/2146), non-modulated = 84.8% (1819/2146).

**Supplementary Fig. 3.**
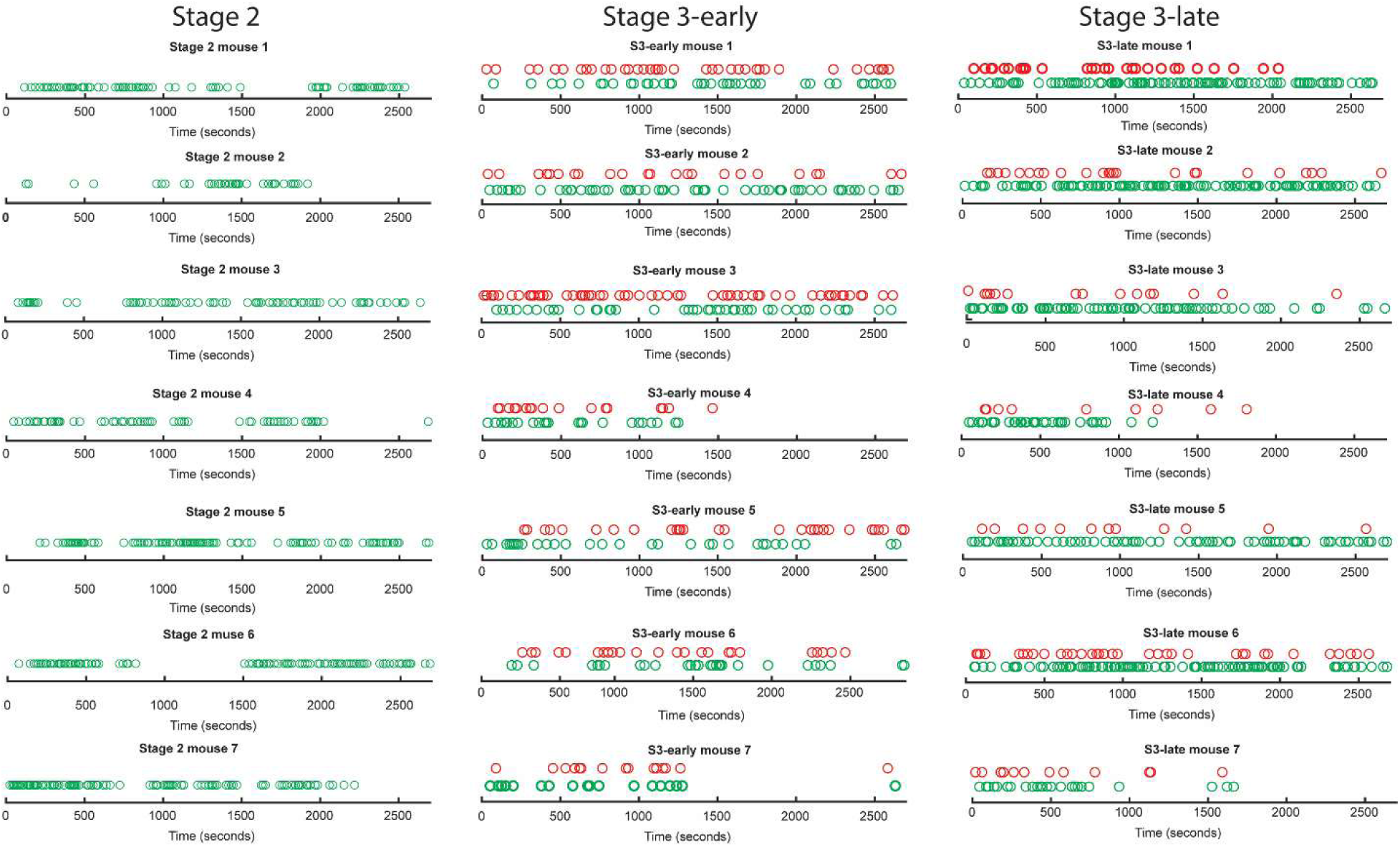
Responses across time during rCPT recording sessions. Distribution of responses (hits-green circles and FAs-red circles) during 45 min (2700 s) recordings sessions.

**Supplementary Fig. 4.**
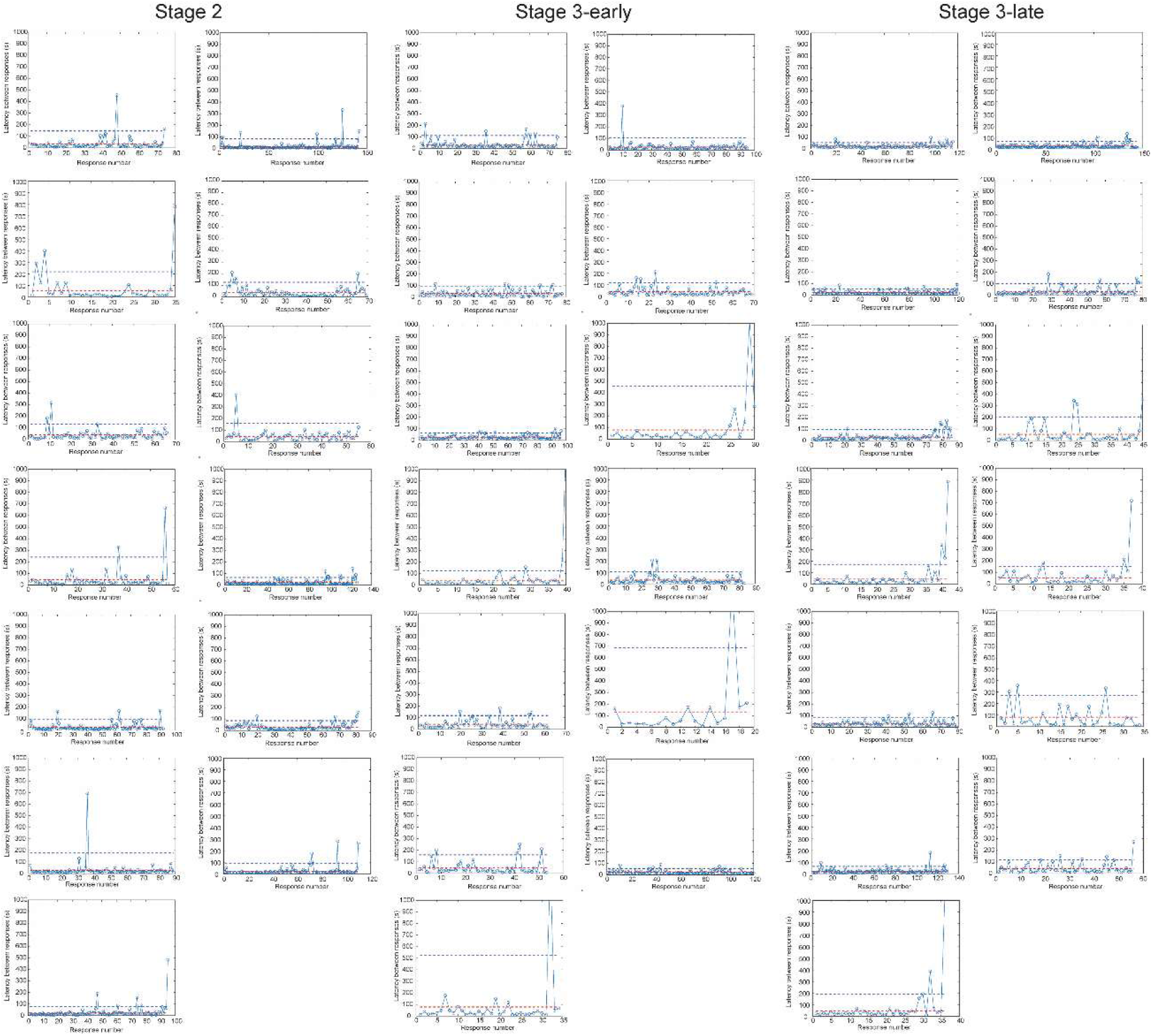
Latency between responses during rCPT recording sessions. Plots showing latency between responses during rCPT recording sessions. Red dotted line shows average latency between two responses for that particular mouse and session. Blue dotted line shows the threshold for disengagement periods (mean + 2DTD) for that particular mouse and session.

**Supplementary Fig. 5.**
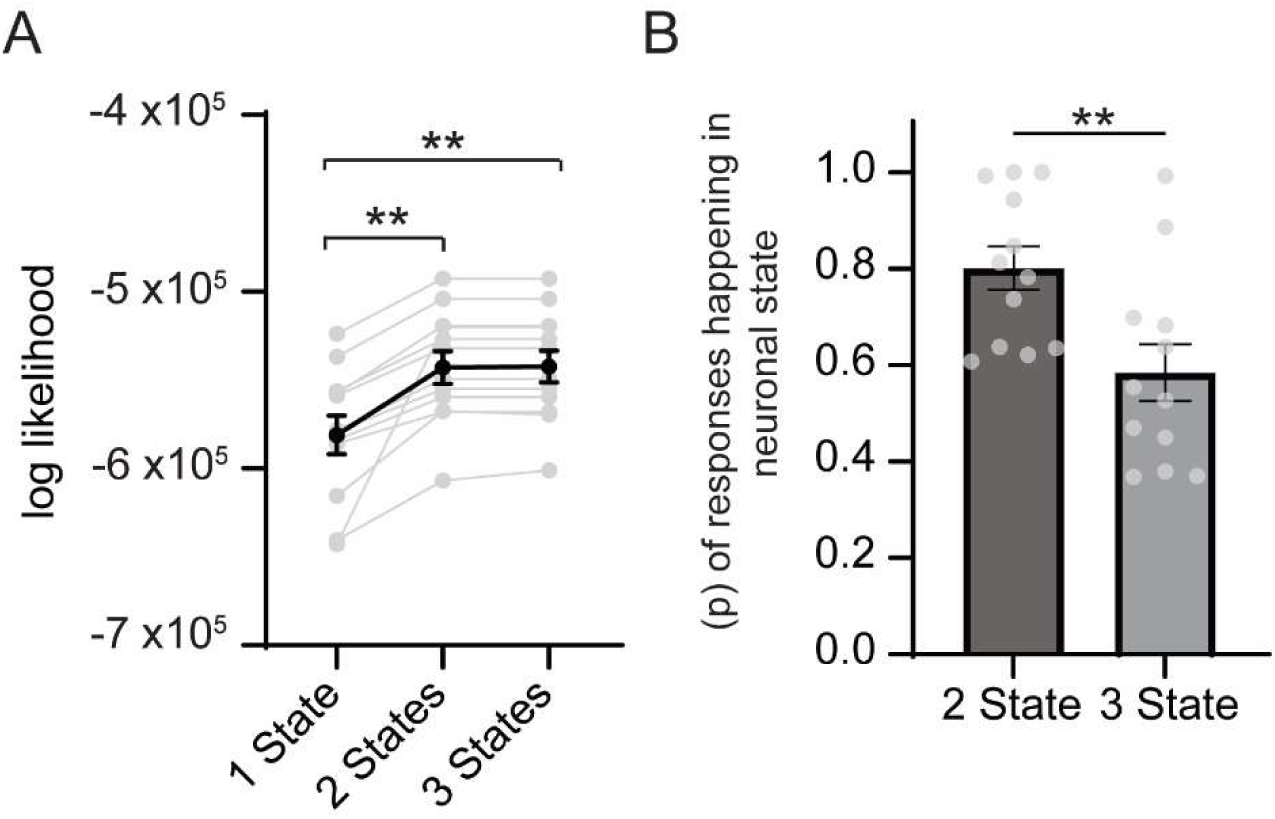
Two-state HMM performance. A) Graph showing the log-likelihood of HMMs fitted with one, two, or three states to neural activity data. Models with two and three states showed significantly higher log-likelihoods compared to the one-state model. No statistical difference was observed between the two- and three-state models. Repeated measures ANOVA F= 23.16 *p* ˂ 0.001 Tuckey’s *posthoc* test 1 state vs 2 states *p*= 0.001, 1 state vs 3 states *p*= 0.001, 2 states vs 3 states *p*= 0.822. B) Bar graph showing the probability of responses occurring in the dominant state for the two- and three-state HMMs. The two-state HMM yielded a higher probability of responses in the dominant state compared to the three-state model (t_22_= 2.94 p= 0.008

**Supplementary Fig. 6.**
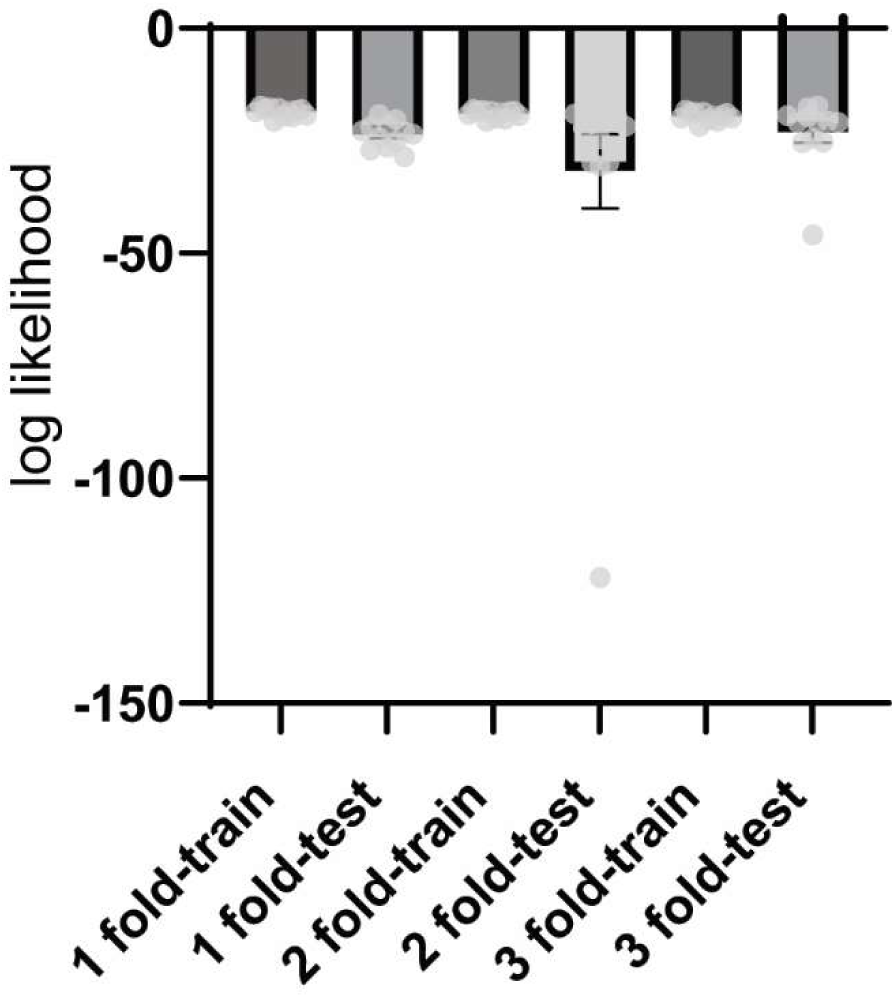
Two-state HMM log-likelihood computed using 3-fold cross-validation within individual behavioral sessions. Sessions were divided into four equal segments, and train versus test sets were defined corresponding to fold number (1-fold; training = first 25% of data, test = second 25% of data). No significant differences in log likelihood across different folds between train and test session. ANOVA F_5,66_ = 1.492 *p* = 0.2043.

**Supplementary Fig. 7.**
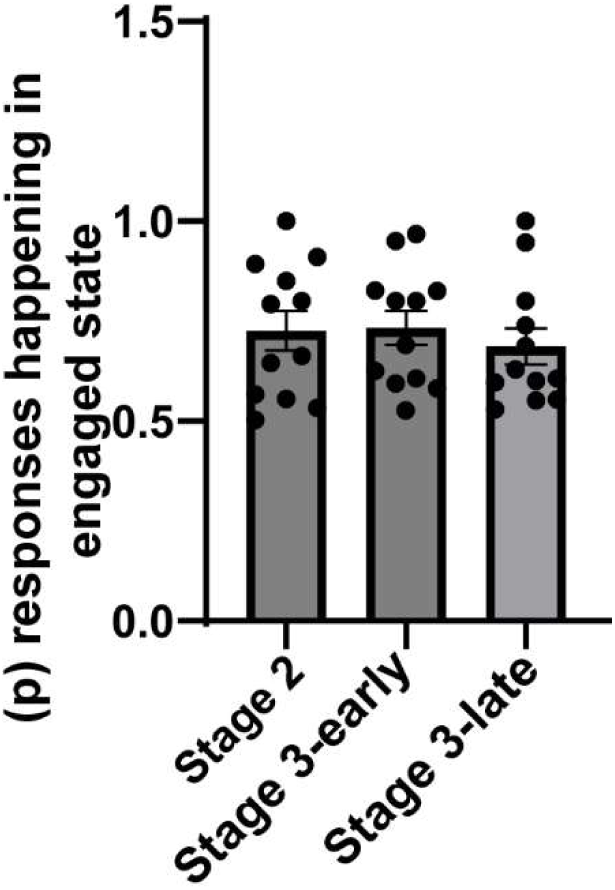
Probability of engaged state across rCPT stages. No statistical difference in the probability of the dominant state given responses between S3-early and S3-late (t_22_= 0.7413 p=0.4663)

**Supplementary Fig. 8.**
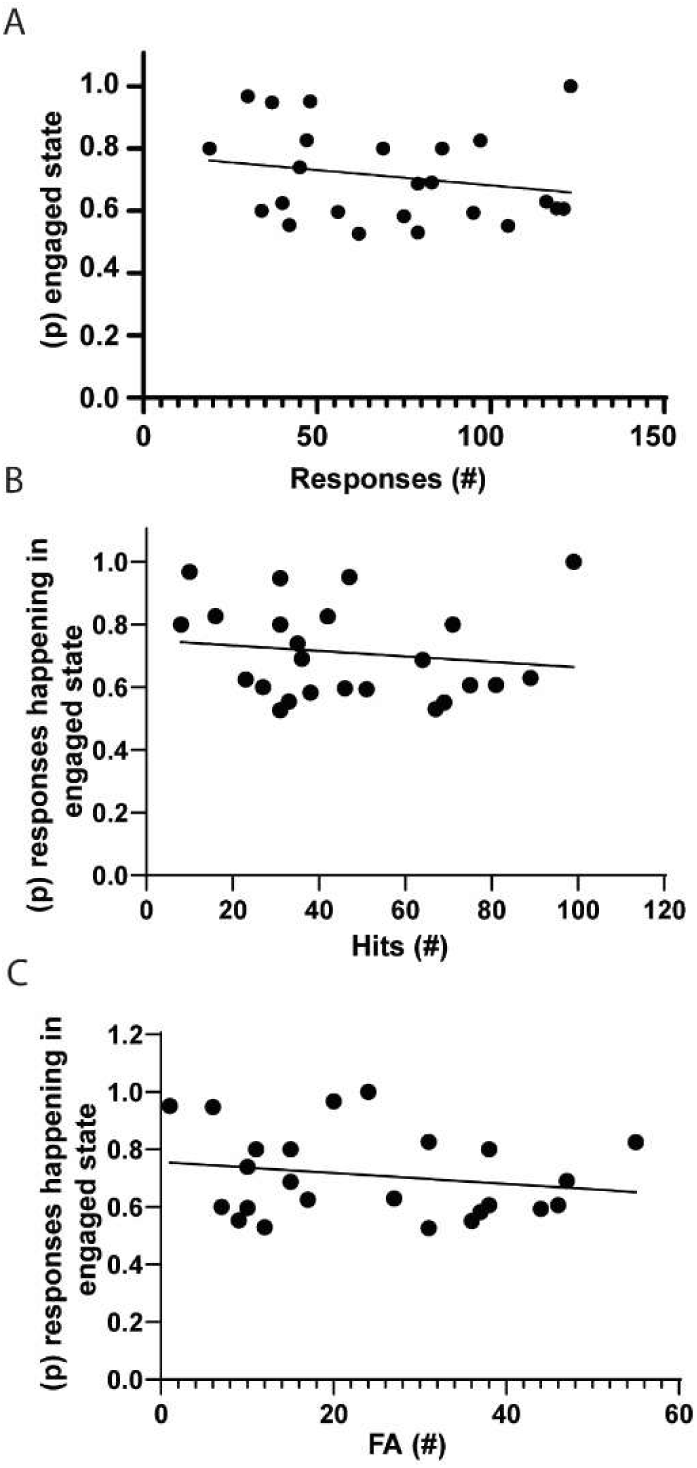
Correlation analysis between the probability for the engaged state against hits and FAs number. A) Correlation graphs showing Bayesian probability of the engaged state given total number of responses. There is no statistical correlation between the number of responses and the probability of responses happening in the engaged state (pearson r= −0.207, p= 0.331). B) Correlation graphs showing Bayesian probability of the engaged state given the number of hits. There is no statistical correlation between the number of hits and the probability of responses happening in the engaged state (pearson r= −0.144, p= 0.502). Correlation graphs showing Bayesian probability of responses happening in the engaged state over the number of FAs. There is no statistical correlation between the number of FAs and the probability of FAs happening in the engaged state (pearson r= −0.194, p= 0.363).

**Supplementary Fig. 9.**
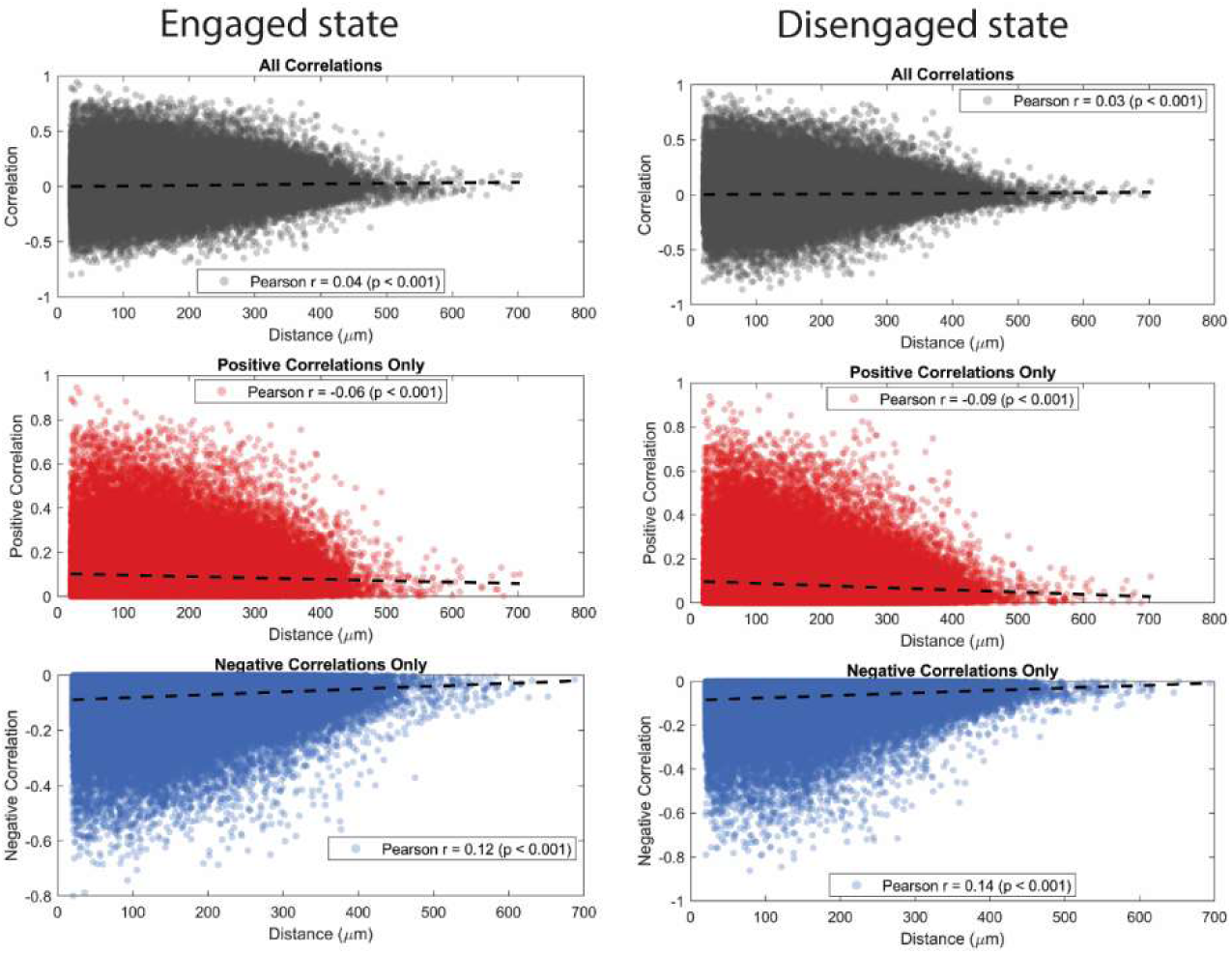
Correlated activity between pairs of PrL neurons decreases with distance across neuronal states associated with task engagement. Analysis of neuronal activity correlation between pairs of neurons and distance during dominant (right) and non-dominant state.

**Supplementary Fig. 10.**
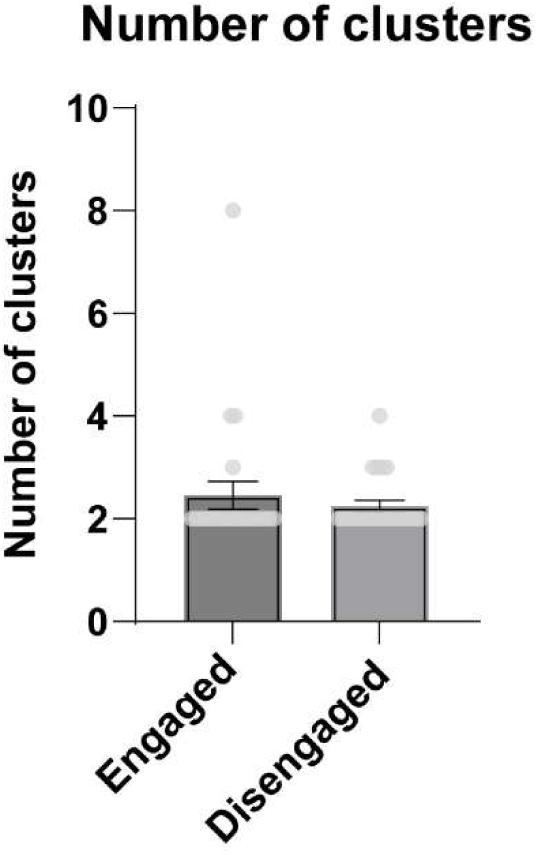
k=2 is the best parameter for k-means cluster analysis during neuronal states across sessions. Bar graph showing the results of Silhouette analysis on correlated activity during rCPT sessions for posterior K-means cluster analysis. There is no significant differences in the number of clusters across engaged and disengaged state (t_23_= 0.7068 p= 0.4868).

**Supplementary Fig. 11.**
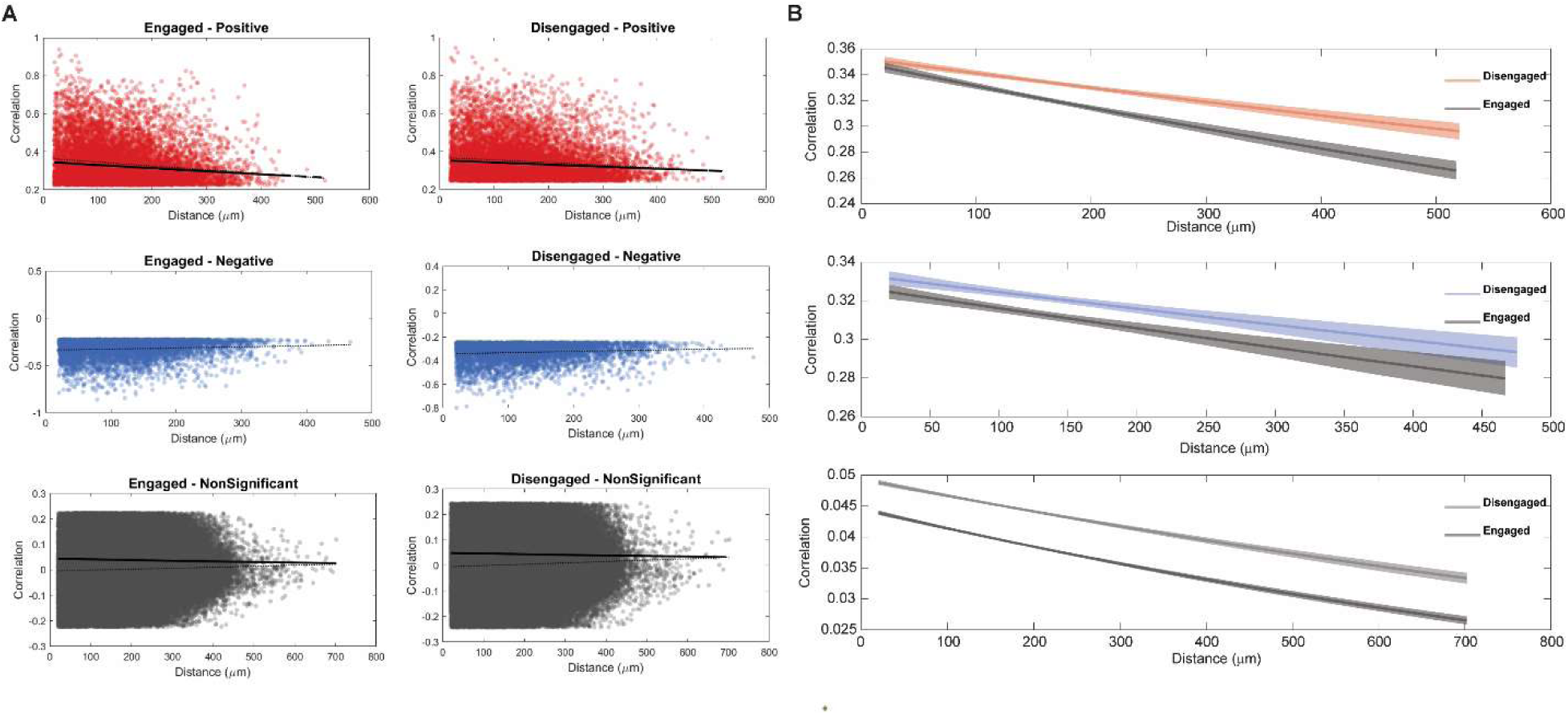
Correlated activity between pairs of PrL neurons decreases with distance across neuronal states associated with task engagement. A) Analysis of neuronal activity correlation between pairs of neurons and distance during (Engaged (right) and Disengaged state divided into significant positive correlated, significant negative-correlated, and non-significant correlated neurons. B) Graphs showing fit of an exponential decay model used to examine correlation strength decline with distance. Decay of positive correlations was significantly steeper in the Engaged state compared to the Disengaged state (*z = 4.21, p*˂ 0.001). Non-significant correlations also showed a significantly steeper decay in the Engaged state (*z = 5.24, p*˂ 0.001). Negative correlations did not differ significantly between states (*z = 1.13, p = 0.257*).

## Acknowledgements

We thank Robert Phillips and other members of the Martinowich and Carr laboratories for helpful comments and suggestions. We also thank Aimee Ormond and Deveren Manley for assistance with animal care.

## Author Contributions

Conceptualization: JMB, HLH, KM Formal analysis: JMB, SA, MST Investigation: JMB, SA, JR, YL Writing-original draft: JMB, KM Writing-review and editing: JMB, SA, JR, YL, MST, GVC, KM Supervision: JMB, GVC, KM Project administration: GVC, KM Funding acquisition: GVC, KM

