## Supplementary figures and images for "Patterns of neural activity in prelimbic cortex neurons correlate with attentional behavior in the rodent continuous performance test"

### Supplementary Fig 1

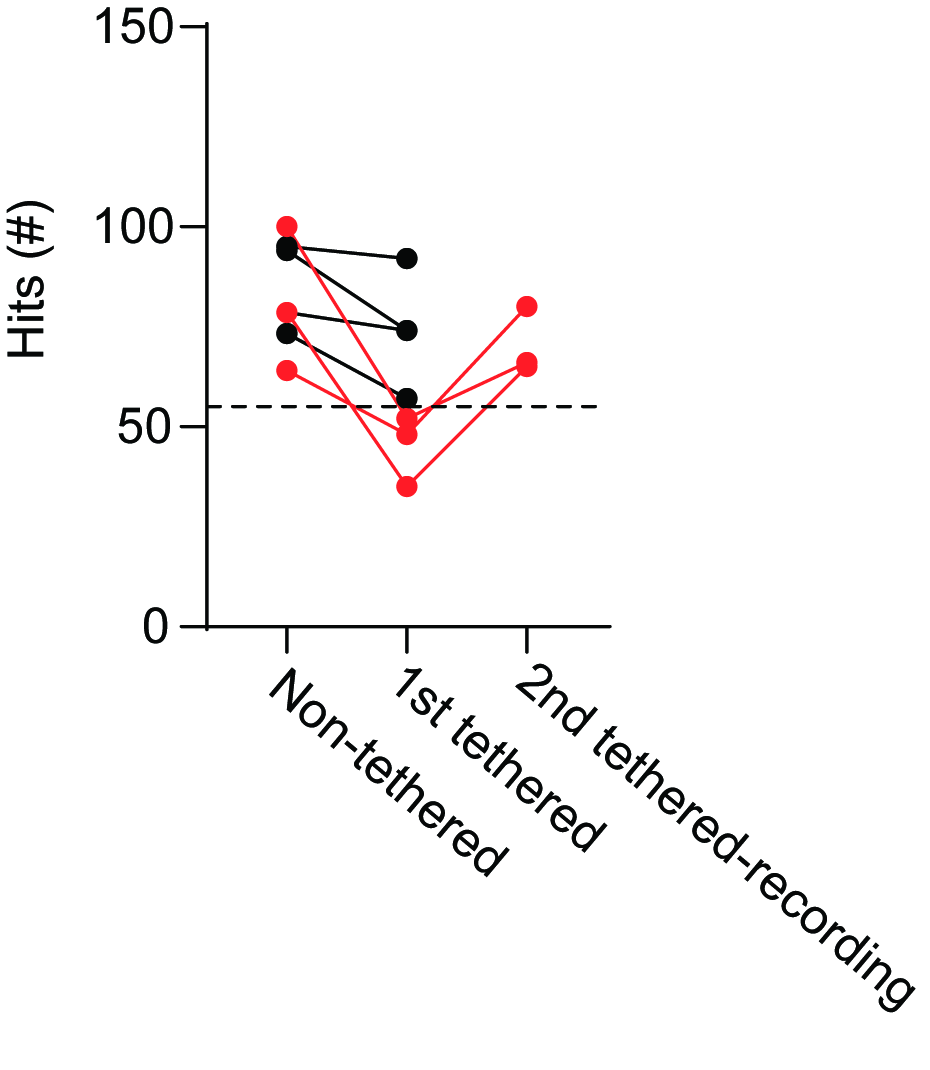

### Supplementary Fig 2

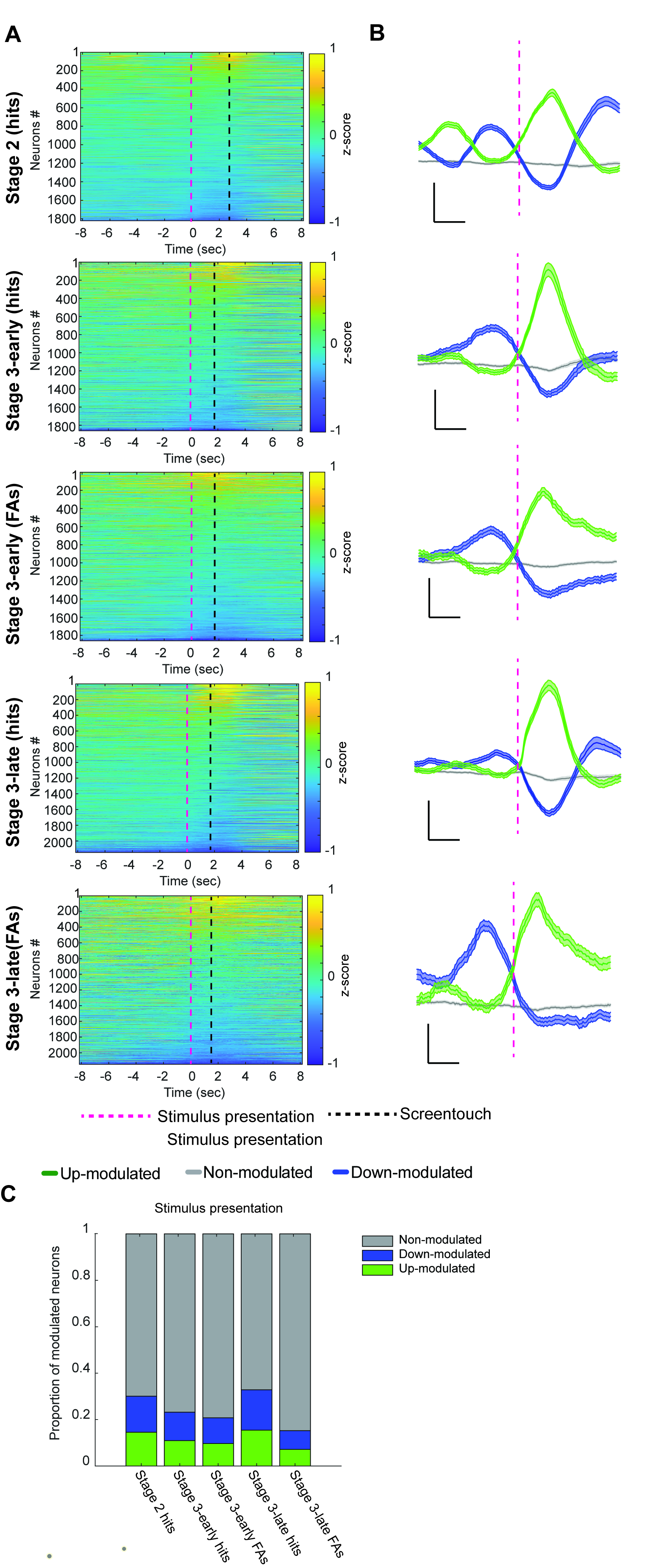

### Supplementary Fig 3

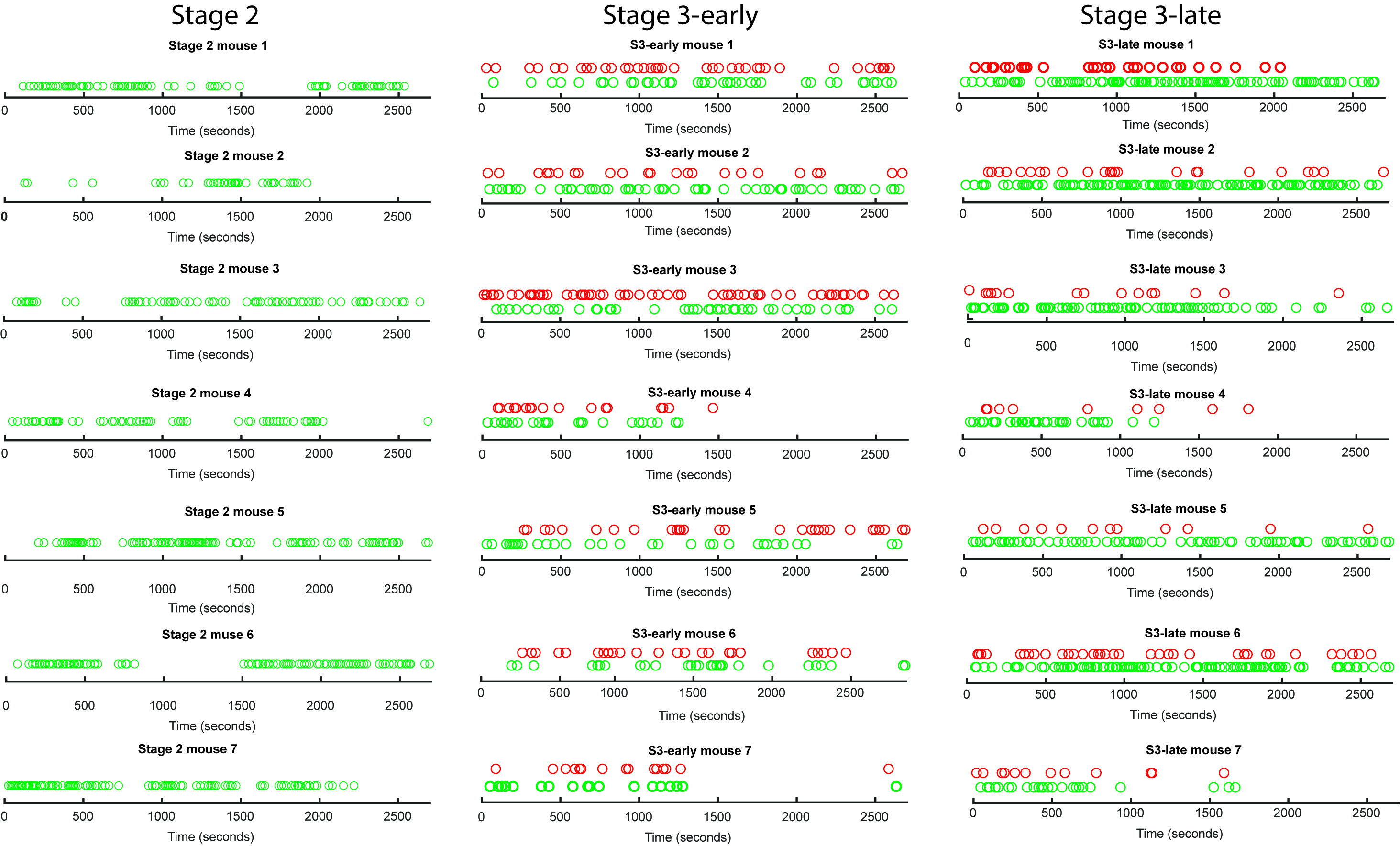

### Supplementary Fig 4

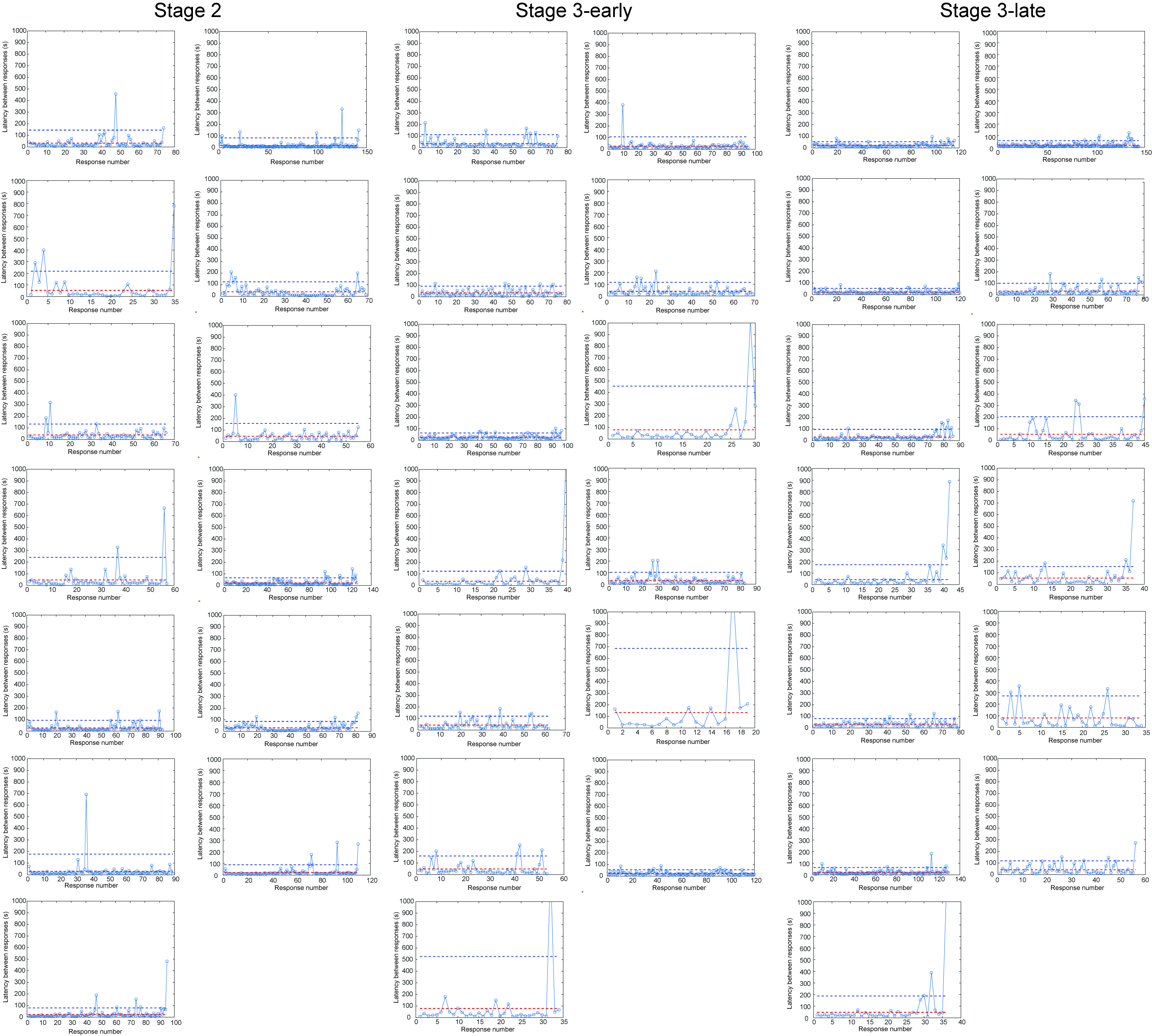

### Supplementary Fig 5

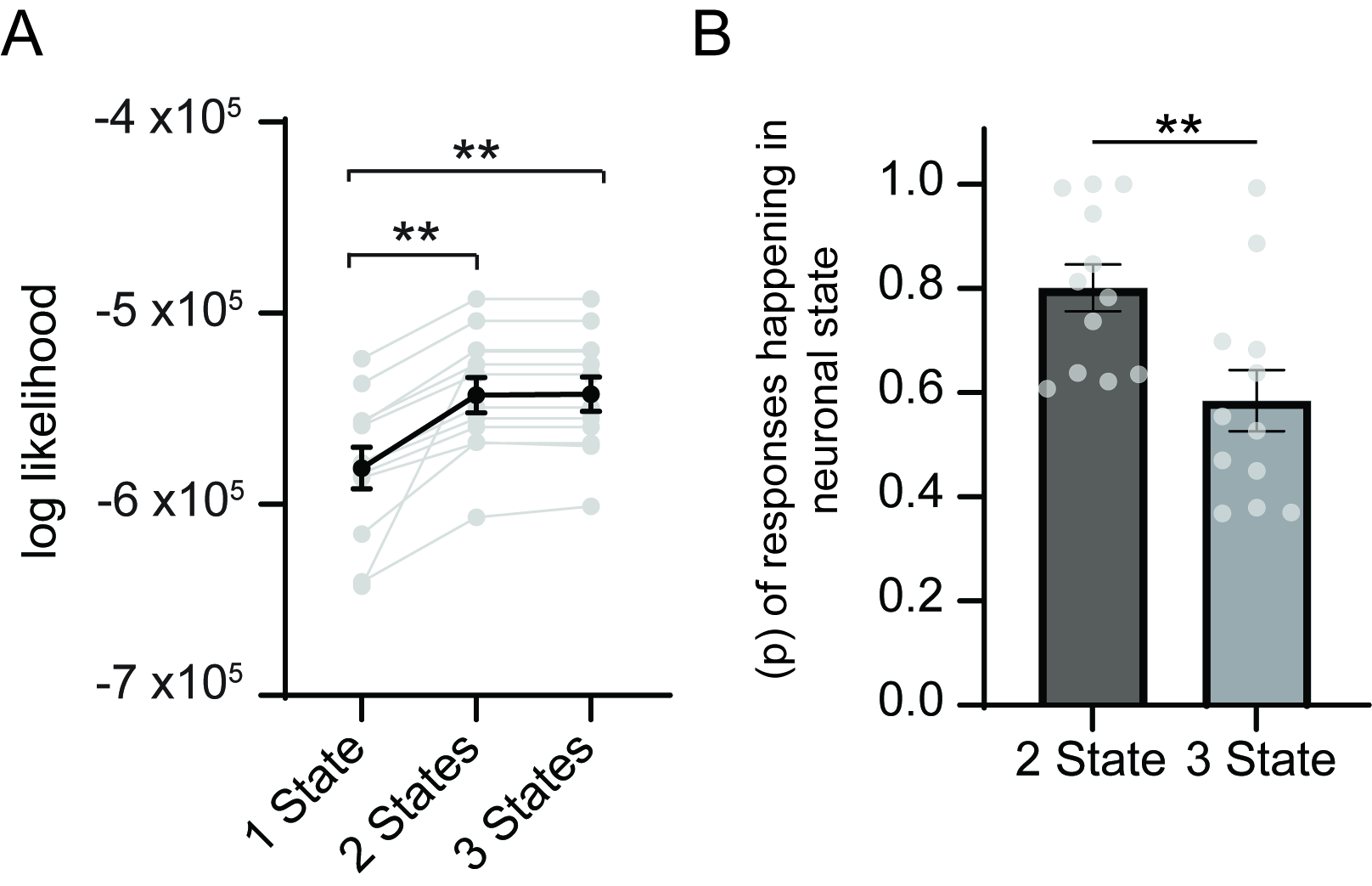

### Supplementary Fig 6

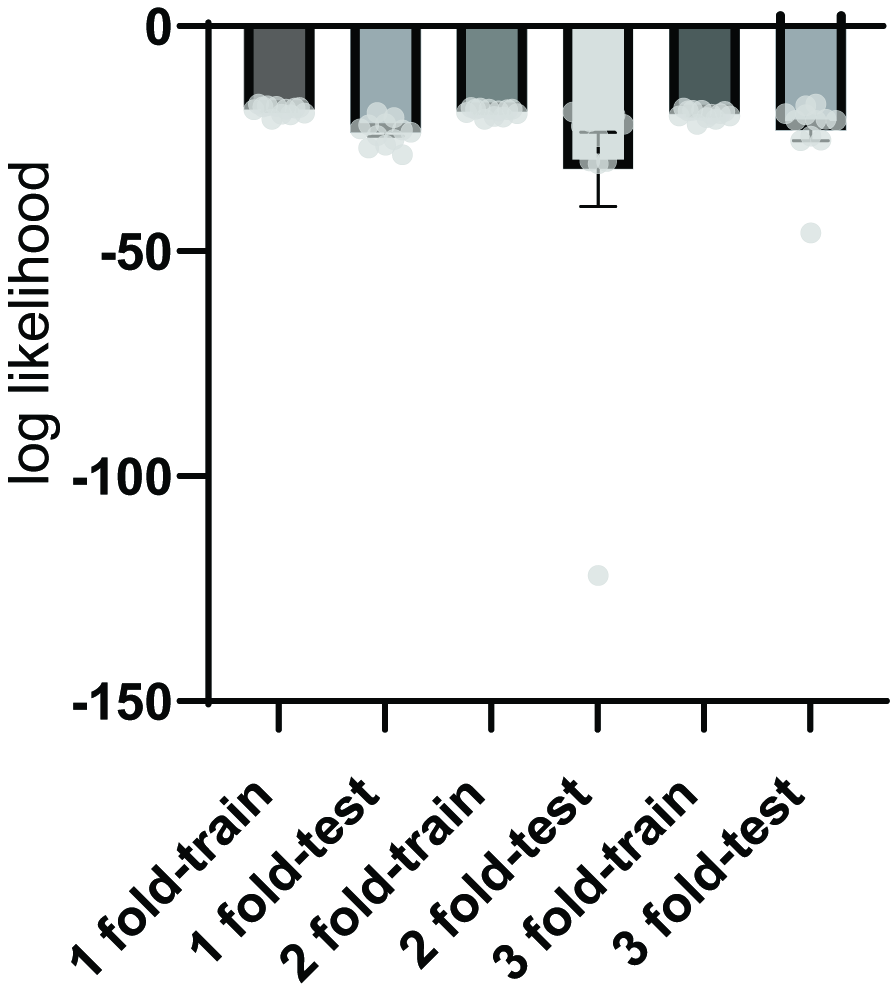

### Supplementary Fig 7

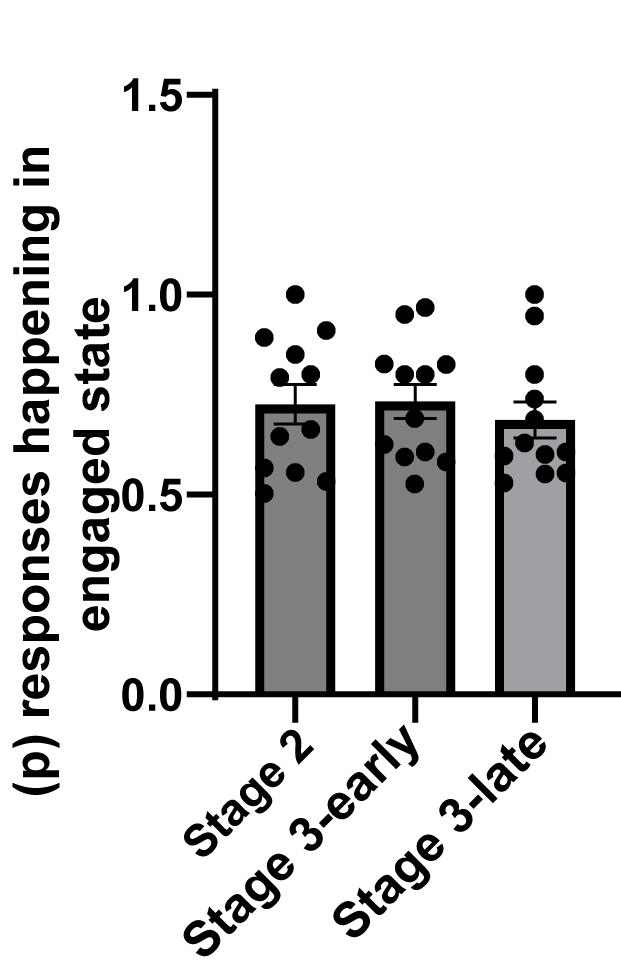

### Supplementary Fig 8

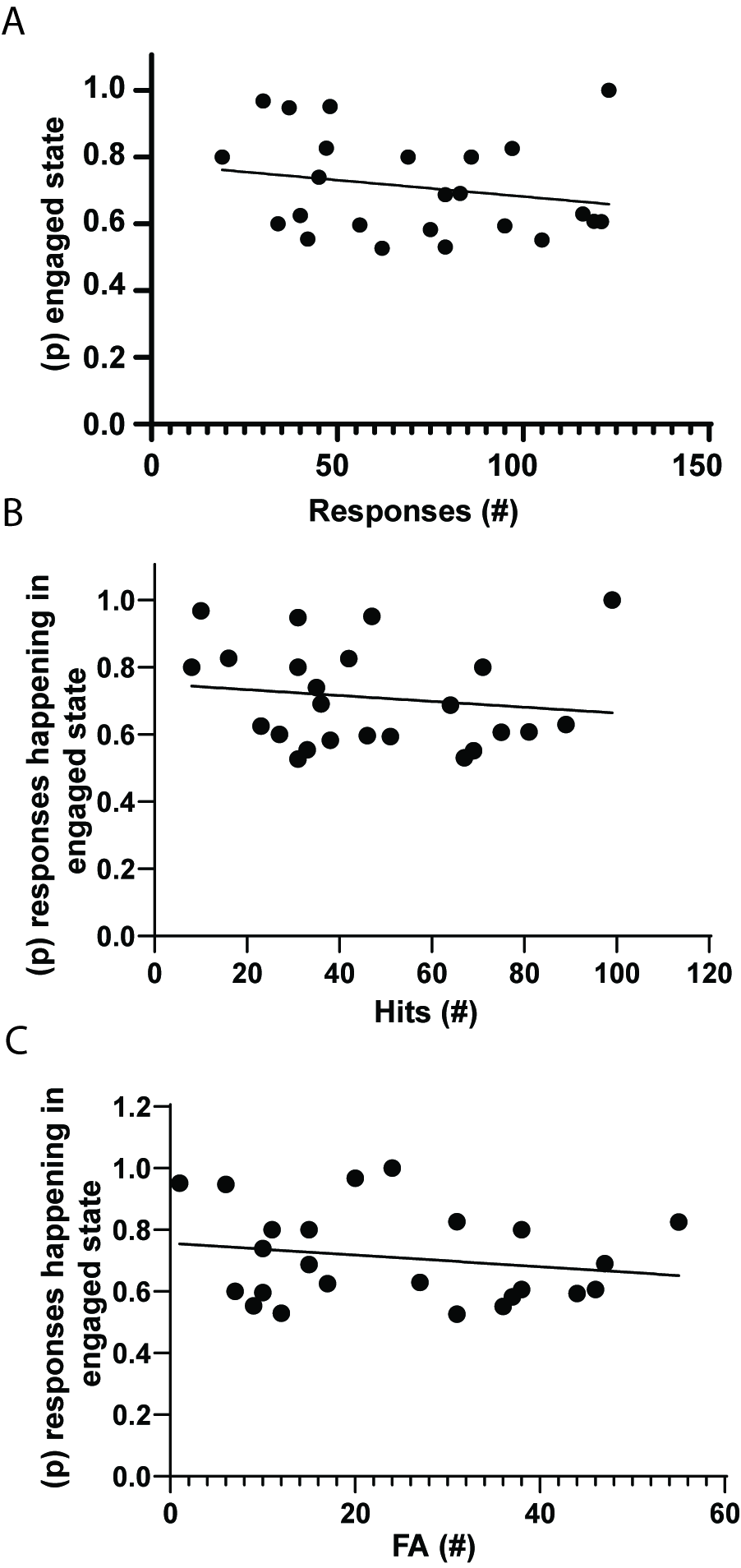

### Supplementary Fig 9

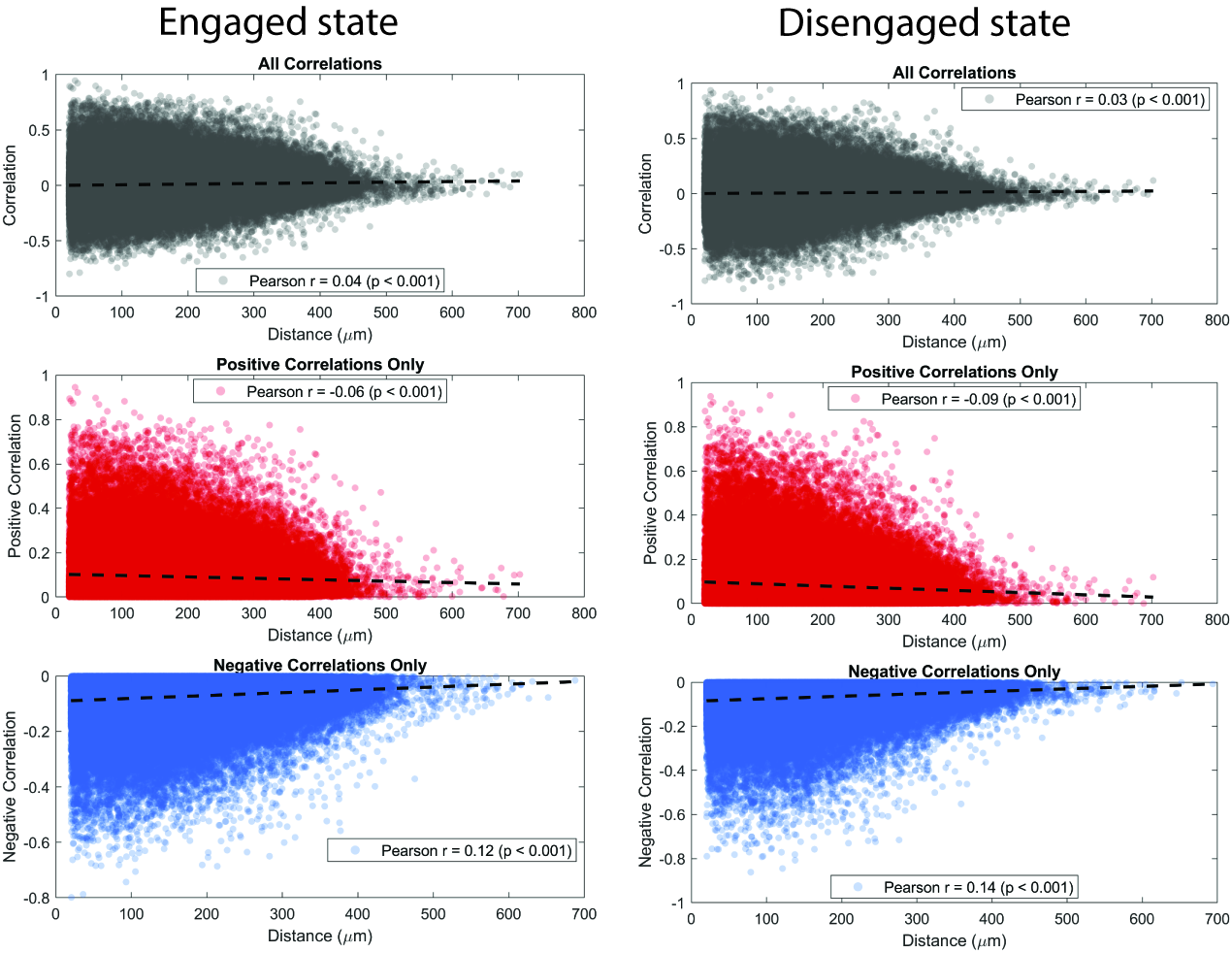

### Supplementary Fig 10

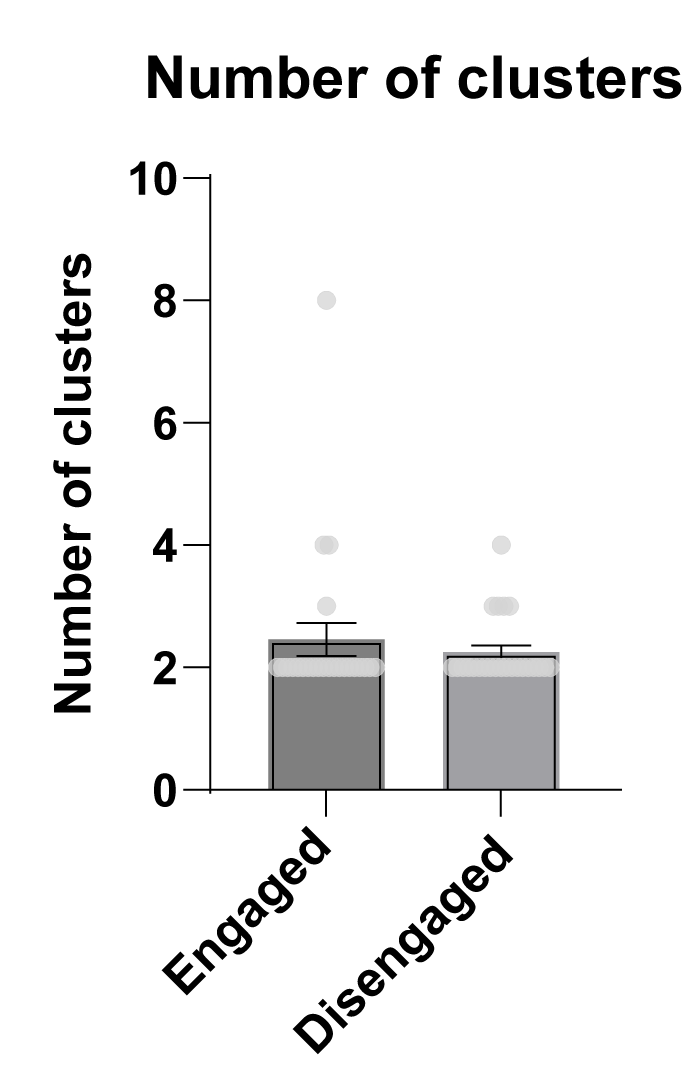

### Supplementary Fig 11

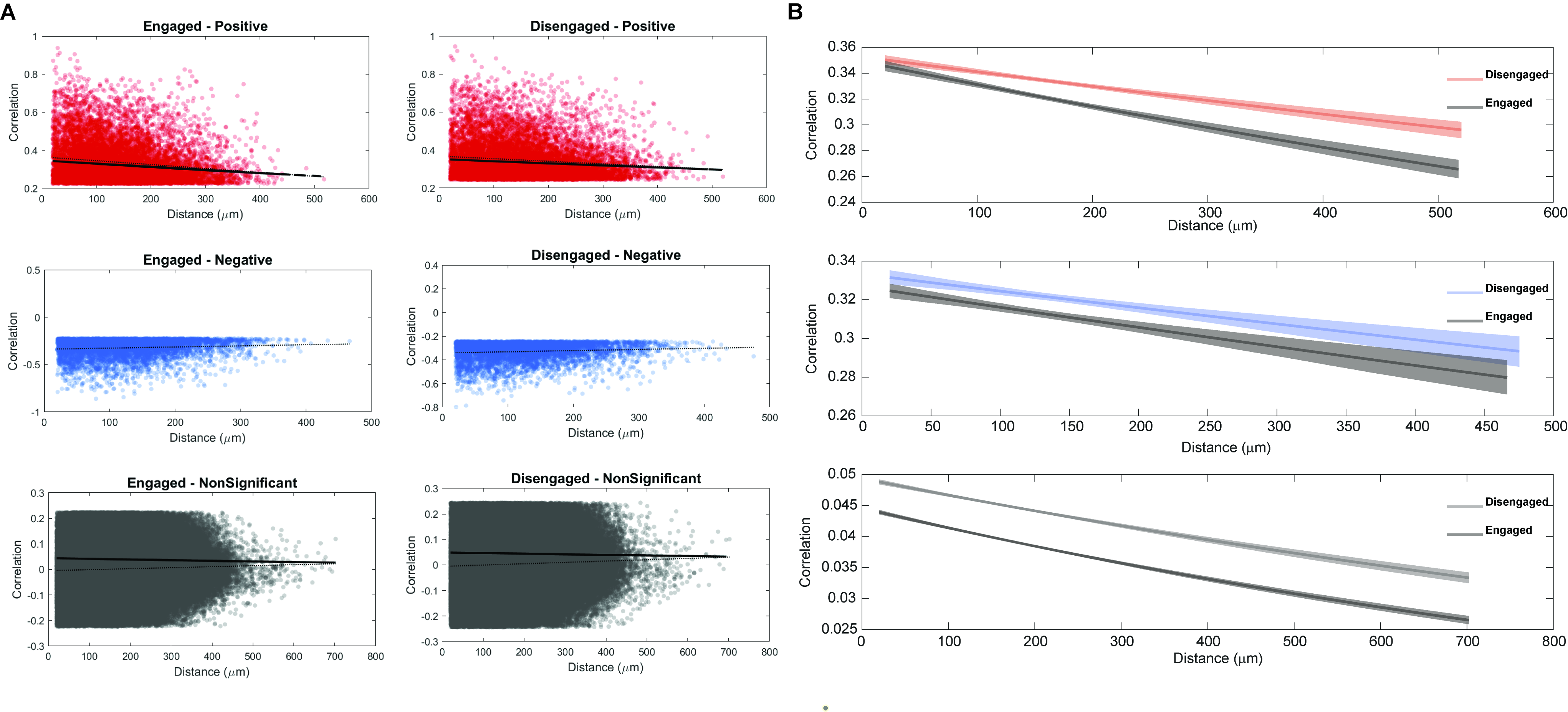
